# Activation of three targets by a TAL effector confers susceptibility to bacterial blight of cotton

**DOI:** 10.1101/2024.06.07.597943

**Authors:** Brendan Mormile, Taran Bauer, Li Wang, Rachel Rivero, Sara C.D. Carpenter, Catherine Danmaigona-Clement, Kevin Cox, Lin Zhang, Xiyu Ma, Terry A. Wheeler, Jane K. Dever, Ping He, Adam J. Bogdanove, Libo Shan

## Abstract

*Xanthomonas* spp. employ transcription activator-like effectors (TALEs) to promote pathogenicity by activating host susceptibility (*S*) genes. Cotton *GhSWEET10* is an *S* gene targeted by a TALE in an early isolate of *Xanthomonas citri* pv. *malvacearum* (*Xcm*), but not by recent field *Xcm* isolates. To understand the pathogenicity shift in *Xcm* and its adaptation to cotton, we assembled the whole genome and the TALE repertoire of three recent *Xcm* Texas field isolates. A newly evolved TALE, Tal7b, activated different *GhSWEET* genes, *GhSWEET14a* and *GhSWEET14b*. Simultaneous activation of *GhSWEET14a* and *GhSWEET14b* resulted in pronounced water-soaked lesions. Transcriptome profiling coupled with TALE-binding element prediction identified a pectin lyase as an additional Tal7b target, quantitatively contributing to *Xcm* virulence alongside *GhSWEET14a/b*. CRISPR-Cas9-based gene editing supported the function of *GhSWEETs* as *S* genes in cotton bacterial blight and the promise of disrupting the TALE-binding site in these genes to control the disease. Collectively, our findings elucidate the rapid evolution of TALEs in *Xanthomonas* field isolates and highlight the virulence mechanism wherein TALEs induce multiple *S* genes simultaneously to promote pathogenicity.

## INTRODUCTION

Cotton (*Gossypium* spp.) stands as a highly valuable crop, cultivated worldwide for its versatility in providing fiber, livestock feed, and oil^1^. The allotetraploid species *G. hirsutum*, commonly known as upland cotton, is predominantly grown for commercial cotton production. Despite its economic importance, cotton yields are threatened by various environmental stresses, including different pathogens. Bacterial blight of cotton (BBC), caused by *Xanthomonas citri* pv. *malvacearum* (*Xcm*), is a destructive foliar disease^2^. The disease can affect any aerial part of the plant during any stage of the bacterial life cycle, and it typically manifests as distinct “water-soaked” lesions. These lesions can progress to become necrotic, causing premature defoliation and boll loss^2,3^. Historically, BBC control relied on measures such as acid-delinted seeds and the cultivation of BBC-resistant cotton varieties^4^. Around 2011, a resurgence of BBC occurred across the southern United States, impacting previously resistant cotton varieties^3,5,6^. The factors contributing to this resurgence, such as pathogen population composition, genomic organization, and virulence dynamics, remain unknown.

The key virulence factors linked to the pathogenicity of *Xanthomonas* spp. are transcription activator-like effectors (TALEs), which emulate the function of eukaryotic transcription factors^7–9^. These TALEs exhibit a highly conserved structure comprising a type III secretion signal at the amino (N)-terminus, followed by a central repeat region (CRR) containing 1.5 to 33.5 tandem repeats, each consisting of nearly identical 33-34 amino acids. The amino acids between different repeats primarily differ at the 12^th^ and 13^th^ residues, designated as the repeat variable di-residue (RVD). Following the CRR, there are nuclear localization signals and an acidic activation domain at the carboxyl (C)-terminus^7–9^. Each RVD within the CRR has a strong selectivity for a specific nucleotide, and the arrangement of RVDs within the CRR corresponds to a binding site within the promoter region of a host target gene, referred to as the effector-binding element (EBE)^4,7^. The first identified TALE, AvrBs3, in *Xanthomonas campestris* pv. *vesicatoria* (*Xcv*), the causal agent of bacterial leaf spot on pepper^10^, induces the expression of *UPA20*, encoding a basic helix-loop-helix (bHLH) transcription factor that regulates plant cell hypertrophy^11^.

The plant *SWEET* gene family, crucial for sugar transport and accumulation in plants^12,13^, encodes proteins primarily located in the cell membrane. These proteins play a pivotal role in the uptake, efflux, and intercellular transport of sugars, with the family divided into four clades based on the specific sugar they transport: clade I and II for hexose, clade III for sucrose, and clade IV for fructose^13,14^. Clade III *SWEET* genes in different plants are commonly targeted by various *Xanthomonas* spp. for pathogenicity, termed susceptibility (*S*) genes. For example, rice *OsSWEET11*^15^, *OsSWEET13*^16^, and *OsSWEET14*^17–19^ are *S* genes targeted by TALEs from *X. oryzae* pv. *oryzae* (*Xoo*). Pepper *UPA16* and cassava *MeSWEET10a* are targeted by TALEs from *Xcv* and *X. axonopodis* pv. *manihotis* (*Xam*), respectively^11,20^. Cotton *GhSWEET10* is targeted by TALE Avrb6 in *Xcm* strain H1005 to induce virulence-associated water-soaking^21^. The upregulation of clade III *SWEET* genes, such as *GhSWEET10* by Avrb6, leads to the efflux of sucrose from the host cell into the apoplast^21,22^. This released sucrose may potentially serve as a carbon and energy source for bacterial infection or alter the apoplast water potential for water-soaked lesion development in plant leaves^16,19,20,23–26^.

*Xcm* H1005 was originally collected from BBC-infected cotton in Oklahoma in 1968^27^. However, reemerging *Xcm* isolates collected from BBC-infected cotton fields in Texas do not induce *GhSWEET10*. Instead, they induce the expression of *GhSWEET14a* and *GhSWEET14b,* two additional members of the clade III *GhSWEET* gene family^21^, highlighting the dynamic nature of *Xcm*-cotton interactions. The shift in the induction of specific *SWEET* genes implies an ongoing evolutionary adaptation, suggesting the potential evolution of new TALEs in *Xcm* to induce alternative *S* genes in cotton. To gain insight into the evolutionary adaptation of *Xcm* and its interaction with cotton, we sequenced and assembled the whole genomes of three recent *Xcm* Texas field isolates and examined their TALE repertoires (TALomes). By generating a series of genetic knockout mutants of individual TALEs and screening them in cotton, a newly evolved TALE, Tal7b, was discovered to contribute to virulence. Tal7b was shown to activate the expression of *GhSWEET14a* and *GhSWEET14b*. While the activation of *GhSWEET14a* or *GhSWEET14b* alone by gene-specific designer TALEs (dTALEs) weakly induced water-soaked lesions, the simultaneous activation of both genes by dTALEs more strongly induced water-soaked lesions, but not completely additively, suggesting partial redundancy of function. CRISPR-Cas9-mediated gene editing targeted at the EBE or coding regions of *GhSWEET10* gene resulted in a loss of susceptibility to *Xcm* H1005, providing additional support for the role of *SWEET* genes in mediating lesion development. To further understand the virulence mechanism and identify additional targets of Tal7b, we deployed a whole-transcriptome analysis along with *in silico* TALE EBE predictions and identified a pectin lyase gene, *GhPL1*, as an additional target of Tal7b. Importantly, *GhPL1* quantitatively contributed to water-soaked lesion development, and when activated alongside *GhSWEET14a* and *GhSWEET14b* together using dTALEs, acted synergistically rather than additively, fully restoring water-soaking to the level induced by Tal7b. Collectively, our findings exemplify the rapid evolution of TALEs in *Xanthomonas* spp. and elucidate a mechanism that the cotton pathogen has evolved to promote disease by simultaneously activating, through a single TALE, members of two functionally distinct gene families that contribute synergistically to water-soaked lesion development.

## RESULTS

### Virulence diversity and whole-genome sequence analysis of recent field *Xcm* isolates

To examine the dynamics of BBC disease, a series of *Xcm* isolates were collected from infected cotton fields across high cotton-yielding regions in Texas between 2010 and 2015, and nine isolates designated *Xcm* TX1 to *Xcm* TX9 were further characterized^21^. Among them, isolates TX4 and TX7 exhibited the most significant upregulation of the *GhSWEET14a* and *GhSWEET14b* genes, whereas isolate TX9 displayed the least or no upregulation of *GhSWEET14a* and *GhSWEET14b*^21^. Given their distinct effects on *GhSWEET* gene expression, these three *Xcm* isolates, TX4, TX7, and TX9, were selected for further investigation in this study (**Figure 1A**). Upon inoculation into the cotton germplasm Acala 44E (Ac44E), *Xcm* isolates TX4 and TX7 elicited robust virulence, characterized by the development of pronounced water-soaked lesions (**Figures 1B and 1C**). This virulence pattern closely resembled that of previously characterized virulent *Xcm* isolates *Xcm* H1005 and *Xcm* MSCT1. In contrast, *Xcm* TX9 showed a very minor or no water-soaked lesion, more similar to *Xcm* HM2.2, a multiple TALE deletion mutant of *Xcm* H1005^28^, which showed nearly no water-soaked lesions upon infection (**Figures 1B and 1C**). Therefore, *Xcm* TX4 and TX7 demonstrate virulence, while *Xcm* TX9 exhibits almost no virulence in cotton Ac44E.

**Figure 1.**
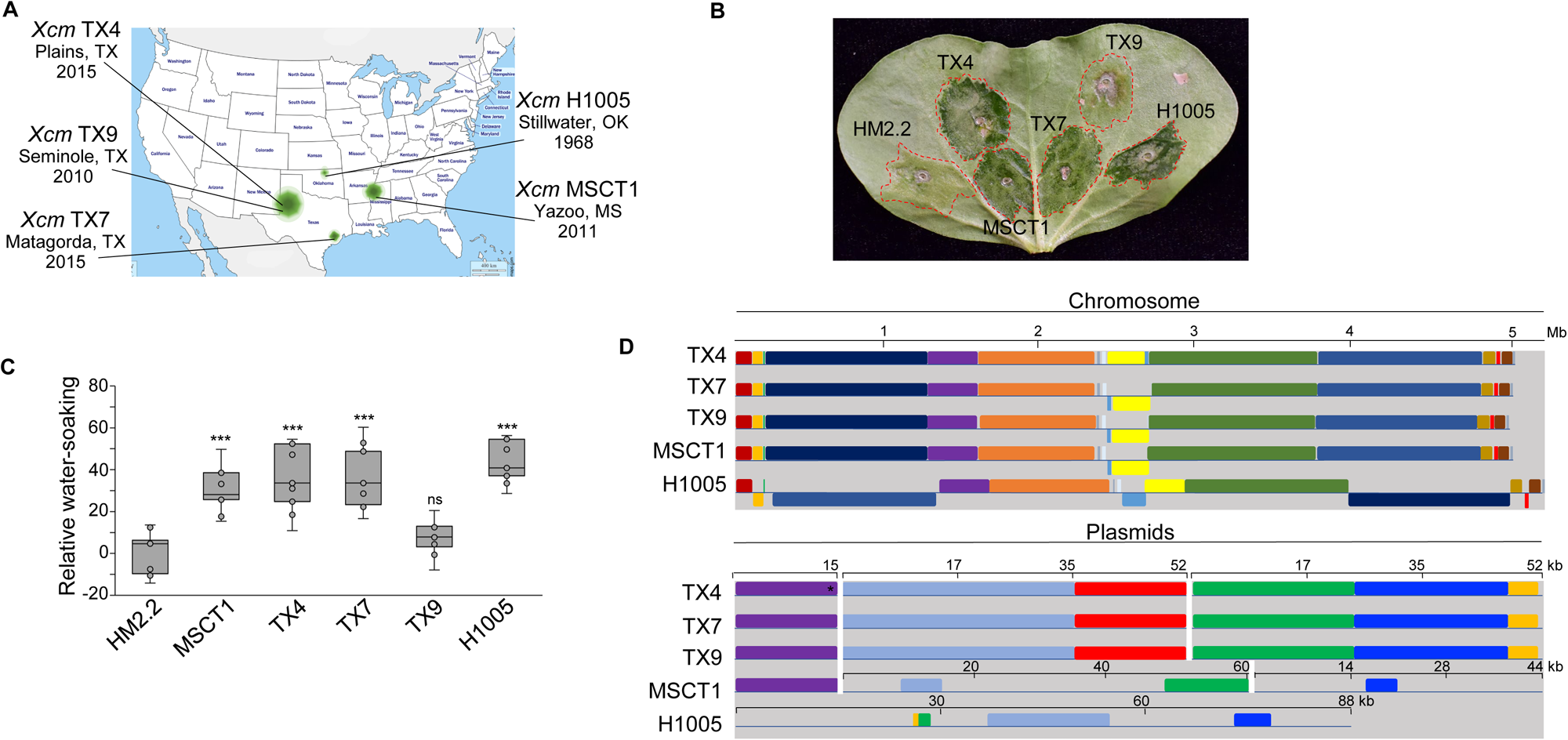
Genomic and virulence comparisons of different *Xcm* isolates. **(A)** Geographical locations and collection years of field *Xcm* isolates used in this study. *Xcm* TX4, TX7, and TX9 were isolated from excised water-soaked lesions on infected cotton leaves and confirmed using Koch’s postulates. *Xcm* H1005 is a derivative of *Xcm* H isolated from Oklahoma in 1968^27^, and *Xcm* MSCT1 was isolated from Mississippi in 2011^30^. **(B)** Individual *Xcm* isolates induce variable degrees of water-soaked lesions in cotton. Different *Xcm* isolates were syringe-inoculated at OD_600_ = 0.1 into ten-day-old cotton cotyledons. Photos were taken five days post-inoculation (dpi). **(C)** *Xcm* TX4 and TX7 elicit more severe water-soaked lesions than *Xcm* TX9. Water-soaked lesions in (**B**) were analyzed using ImageJ FIJI following the established protocol^75^. The data are shown as the negative grayscale means of water-soaked lesions for *Xcm* isolates relative to *Xcm* HM2.2 used as a control. The grayscale mean was determined by averaging the grayscale values in the RGB channels from ≥ 9 independent repeats. The asterisks indicate a significant difference (*** *p* < 0.001, ns = no significance) compared to *Xcm* HM2.2 control using a two-tailed Student’s *t*-test. **(D)** Genome comparisons of different *Xcm* isolates. Genome alignment of *Xcm* isolates was conducted using progressiveMAUVE^66^ with default parameters. Colored rectangles represent locally collinear blocks of homology without any rearrangement across the aligned sequences. *Xcm* TX4, TX7, and TX9 each harbor a circular 5-megabase pair (Mb) chromosome, two 52-kilobase plasmids (*pXCM52.4kb* and *pXCM52kb*), and a 15-kb plasmid (*pXCM15kb*), with overall sequence identities exceeding 99% among the three isolates. *Xcm* TX4 contains an inverted region (colored in yellow) in the chromosome and an additional tandem 12-bp (*) sequence in *pXCM15kb* compared to *Xcm* TX7 and TX9. Mississippi isolate *Xcm* MSCT1 harbors a circular 5-megabase pair chromosome, one 15-kb plasmid (*pMSCT15kb*), one 60-kb plasmid (*pMSCT60kb*), and one 44-kb plasmid (*pMSCT44kb*). *Xcm* H1005 contains a 5-megabase pair chromosome and one 88-kb plasmid. The genome sequences of *Xcm* H1005 and *Xcm* MSCT1 were retrieved from the National Center of Biotechnology Information (NCBI) database. The experiments in B and C were repeated four times with a similar result.

To explore potential genome variations contributing to virulence, we employed whole-genome sequencing with the Pacific Biosciences (PacBio) single-molecule real-time (SMRT) sequencing platform, generating long reads capable of spanning the complete length of the repeat-rich TALE genes for efficient *de novo* genome assembly. The resulting assemblies revealed a common genomic architecture among Texas *Xcm* isolates, each harboring a 5-megabase (Mb) circular chromosome, along with one 15-kilobase^29^ plasmid *(pXCM15kb*) and two 52-kb plasmids (*pXCM52.4kb* and *pXCM52kb*) (**Figure 1D, Supplementary Figure 1, and Supplementary Tables 1 and 2**). Genome alignment demonstrated remarkable similarities in overall genomic structures among *Xcm* TX4, TX7, and TX9 (**Figure 1D and Supplementary Figure 1**). Noteworthy differences emerged as an inverted region within the chromosome and an additional tandem 12-bp segment in plasmid *pXCM15kb* of *Xcm* TX4 compared to TX7 and TX9 (**Figure 1D**). Despite variations in virulence, these findings suggest that *Xcm* TX4, TX7, and TX9 share an almost identical overall genome sequence, indicating a likely clonal relationship.

*Xcm* MSCT1 was isolated from a bacterial blight-infected cotton field in Mississippi during the 2011 outbreak^30^. To facilitate a comparative analysis of *Xcm* isolates associated with the BBC resurgence in different states, genome alignment was performed to compare Mississippi isolate MSCT1 with Texas isolates. Given the high genomic similarity among Texas *Xcm* isolates, TX4 was chosen as a representative for the pairwise comparison. MSCT1 harbors a 5-Mb circular chromosome, one 15-kb plasmid (*pMSCT15kb*), one 60-kb plasmid (*pMSCT60kb*), and one 44-kb plasmid (*pMSCT44kb*) (**Figure 1D**). In line with different Texas isolates, the chromosome and plasmid *pMSCT15kb* of *Xcm* MSCT1 are highly similar to those of *Xcm* TX4. Notably, dissimilarities emerge in the two large plasmids, with relatively few locally collinear blocks (**Figure 1D**).

However, *Xcm* H1005, an early isolate collected in 1968 from Oklahoma^27^, exhibited high divergence from *Xcm* TX4 and MSCT1 in both the chromosome and plasmids. Substantial genomic alternations, including chromosome translocations, inversions, and deletions, were observed in the chromosome between *Xcm* TX4 and H1005. The divergence is even more pronounced in the plasmids, where *Xcm* H1005 carries only one plasmid, which has very limited collinear blocks with the 52-kb plasmids of *Xcm* TX4. Apparently, both Mississippi isolate MSCT1 and Texas *Xcm* isolates are highly divergent from the early Oklahoma isolate (**Figure 1D**).

### Diversity in the TALE repertoire among different *Xcm* isolates

TALEs are key determinants for *Xanthomas* pathogenicity^7–9^. To gain a better understanding of the TALE diversity and evolution, we analyzed the TALE repertoires of Texas *Xcm* isolates using the AnnoTALE tool^31^. Each Texas *Xcm* isolate has a total of eight intact TALE genes: one on the chromosome (Tal1), five on plasmid *pXCM52.4kb* (Tal2 to Tal6), and two on plasmid *pXCM52kb* (Tal7b and Tal8) (**Figure 2A**). Of note, we named the TALEs in Texas isolates based on their order in the genome from the origin of replication and by replicon (chromosome first, then plasmids in order)^32,33^. The nomenclature of “Tal7b” is attributed to the presence of a partial TALE sequence immediately upstream, with an incomplete 5’ region and only one repeat. *Xcm* TX4 and MSCT1 exhibited a similar TALE composition and identity across their genomes (**Figures 2A, 2B, 2C, and 2D**). Note that the TALEs of MSCT1 were previously named numerically as well^30^, but not in order of position in the genome, so the numbers do not correspond to those of the Texas isolate TALEs. Immunoblot analysis with α-TALE antibodies revealed that the shared number and relative sizes of TALEs in *Xcm* MSCT1 and the Texas isolates are not conserved in strain H1005 and strain N1003, a derivative of *Xcm* N collected from Burkina Faso in 1968^27^ (**Figure 2B)**.

**Figure 2.**
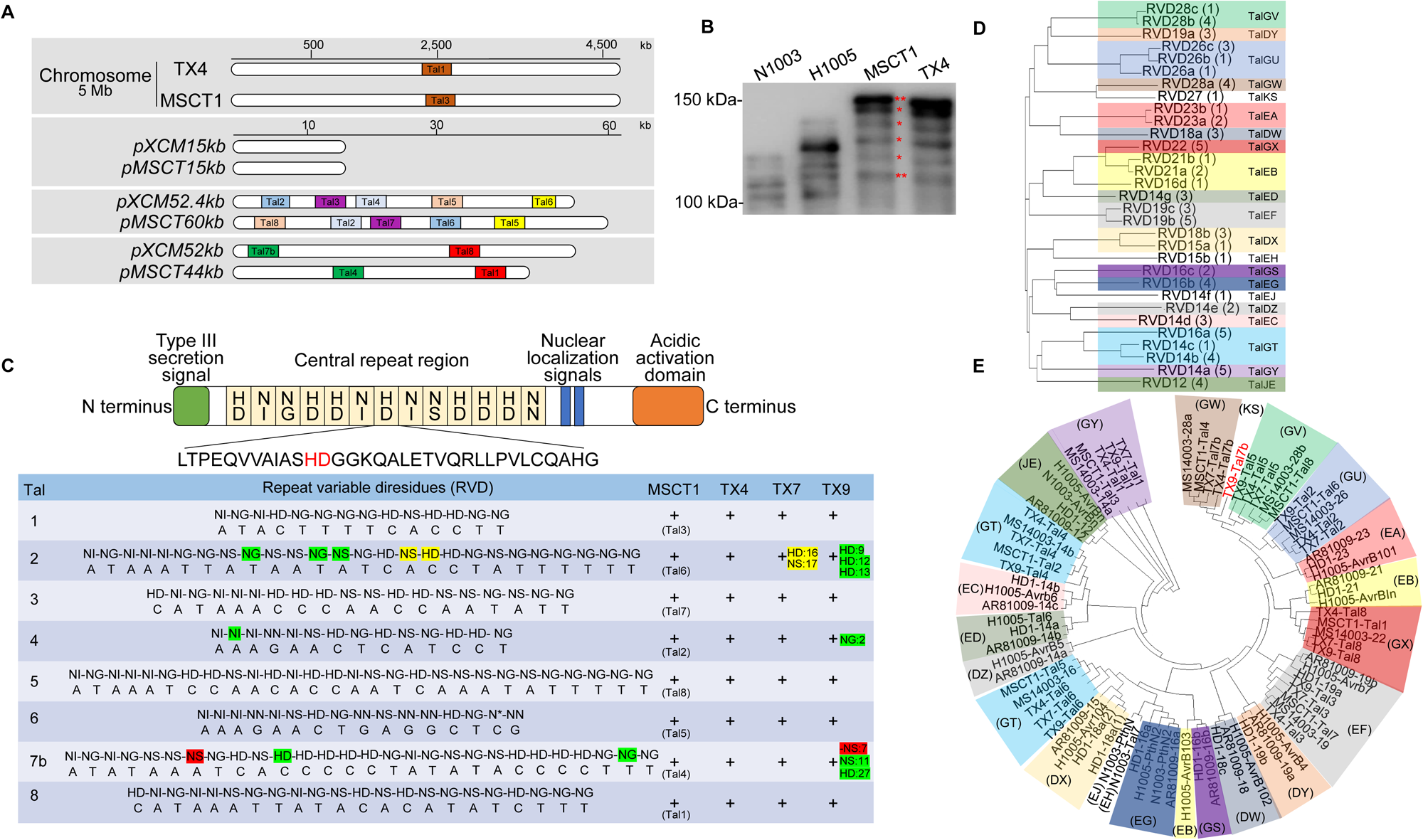
Identification and comparison of TALE repertoires in reemerging *Xcm* isolates and their phylogenetic analysis. **(A)** TALE gene distribution in the chromosome and plasmids of *Xcm* TX4 and MSCT1. TALEs in *Xcm* TX4 were designated numerically from the origin of replication in the chromosome and plasmids. TALEs in *Xcm* MSCT1 were designated previously using a different scheme^30^. The same-colored boxes indicate TALEs with identical RVD sequences. **(B)** *Xcm* TX4 and MSCT1 share similar TALome profiles distinct from those of *Xcm* H1005 and N1003. *Xcm* strains were cultured in nutrient broth and cell homogenates were subjected to immunoblot analysis using α-TALE antiserum^71^. Molecular weight (kDa) is shown on the left of the gel. Asterisks (*) indicate annotated TALEs at their predicted sizes. The experiment was repeated twice with a similar result. **(C)** The repeat variable di-residues (RVD) and predicted effector binding elements (EBE) of the TALEs from *Xcm* MSCT1 and TX4, TX7, and TX9. RVD sequences are identical within the TALome of *Xcm* TX4 and MSCT1. *Xcm* TX9 exhibits three RVD mismatches in Tal2, one mismatch in Tal4, and two mismatches and a deletion in Tal7b relative to *Xcm* TX4 and MSCT1. *Xcm* TX7 has two RVD mismatches within Tal2 relative to *Xcm* TX4 and MSCT1. The names of the TALEs from *Xcm* MSCT1 are shown in parentheses. **(D)** Relationships of TALE from various *Xcm* isolates based on the sequences of their predicted DNA targets, made using FuncTAL^35^. Thirty-one unique RVD sequences from eighty TALEs, obtained from nine *Xcm* isolates (detailed in Results) with available full genome sequences, were analyzed. FuncTAL clusters TALEs based on functional relatedness by calculating similarities in predicted EBE probabilities according to the RVD-DNA code. Highlighted boxes with different colors show the different classes (Tal followed by a two-letter code, such as TalGV) assigned by AnnoTALE. Individual TALEs in the tree are designated as “RVD” followed by the number of repeats within their central repeat region (CRR) and a lowercase letter to distinguish among TALEs with the same number of repeats. Numbers in parentheses indicate TALEs from different isolates with the same RVD sequence. The genome sequences of *Xcm* HD-1, AR81009, and MS14003 were retrieved from the NCBI database. **(E)** Phylogenetic analysis of TALEs from various *Xcm* isolates. Eighty TALEs in (**D**) were analyzed using DisTAL^35^ which clusters based on CRR nucleotide sequence similarities. Highlighted boxes with different colors represent different TALE classes according to AnnoTALE, with the two-letter designations in parentheses, as in (**D**). Only TALEs from *Xcm* isolates with available whole-genome sequences were used in this analysis.

The CRRs harboring RVDs in TALEs of Texas *Xcm* isolates and *Xcm* MSCT1 range in length from 14 to 28 copies of the tandem amino acid sequence. Given the importance of RVD composition in the recognition and binding of TALEs to their target DNA sequence, we compared TALE RVDs among recent *Xcm* isolates: MSCT1, TX4, TX7, and TX9. Notably, water-soaked lesion-inducing *Xcm* isolates MSCT1, TX4, and TX7 share nearly identical RVD sequences within all eight TALEs, except Tal2_TX7_, which contains two RVD differences relative to its orthologs in *Xcm* MSCT1 and TX4 (**Figure 2C**). In contrast, the virulence-attenuated *Xcm* TX9 exhibits several RVD dissimilarities in Tal2, Tal4, and Tal7b compared to the counterparts in *Xcm* MSCT1 and TX4 (**Figure 2C**). Considering the important role of the sequence of RVDs for directing the TALE protein to its host target gene and inducing susceptibility, the divergence of these sequences in Tal2, Tal4, and Tal7b of the weakly virulent strain suggests that one or more of these TALEs may contribute to bacterial pathogenicity and water-soaked lesion development.

Next, we conducted an analysis to assess the functional relatedness of TALEs derived from the nine *Xcm* strains collected across diverse geographic locations and spanning different isolation dates (**Supplementary Table 3**). In addition to *Xcm* TX4, TX7, TX9, MSCT1, N1003, and H1005, we included three other *Xcm* strains with available full genome sequencing data at the time of our analysis: MS14003 isolated from Mississippi in 2014, AR81009 isolated from Argentina in 1981^34^, and HD-1 isolated from China in 2017 (https://www.ncbi.nlm.nih.gov/datasets/genome/GCF_009671025.1/, GenBank ID: GCA_009671025.1). In total, 80 TALEs are present across nine *Xcm* strains. Among the 80 TALEs, thirty-one unique RVD sequences are represented, with the number of repeats in individual proteins ranging from 12 to 28 (**Supplementary Figure 2**). These 31 RVD sequences were grouped into 21 classes based on their predicted corresponding DNA target sequences (EBEs), using FuncTALE^35^ (**Figure 2D**). Since TALEs can tolerate a certain number of mismatches between their RVDs and corresponding EBEs, the relatively high number of EBE classes compared to unique RVDs suggests a greater degree of sequence diversity within the CRRs. This plasticity is possibly driven by frequent recombination events within the CRRs, allowing for adaptation to different host genotypes^36^. Further analysis, using DisTAL^35^ revealed that the groups based on predicted EBEs almost exclusively consist of phylogenetically related TALEs (**Figure 2E**). For example, the FuncTAL class TalGY contains Tal1 from three Texas isolates and the corresponding TALE in two Mississippi isolates. Moreover, phylogenetically related reemerging *Xcm* isolates exhibited similar TALomes, indicating a common ancestral origin. Notably, despite the RVD polymorphisms observed in Tal2 and Tal4 among Texas isolates (**Figure 2C**), Tal2 and Tal4 from TX9 belong to the same FuncTAL class as Tal2 and Tal4, respectively, from *Xcm* TX4 and TX7 (**Figure 2E**). This indicates that differences in TAL2 and TAL4 RVD composition in different Texas isolates do not result in gross alterations to their EBE target sites. In contrast, Tal7b from *Xcm* TX9 belongs to a distinct FuncTAL class compared to Tal7b from *Xcm* TX4 and TX7, implying a different EBE target site for Tal7b from *Xcm* TX9 in comparison to *Xcm* TX4 and TX7 (**Figure 2E**). This also hints at a critical role of Tal7b in mediating bacterial virulence.

### Tal7b mediates *GhSWEET14a/b* induction and water-soaked lesion development

Texas *Xcm* isolates TX4 and TX7 induced the expression of *GhSWEET14a* and *GhSWEET14b* in cotton^21^. To identify the TALE responsible for this induction, we constructed a broad-host-range vector *pKEB1* expressing individual TALEs from *Xcm* TX4 and transformed them into *Xcm* HM2.2, a multiple TALE deletion mutant of *Xcm* H1005. Quantitative reverse transcription polymerase chain reaction (RT-qPCR) was used to analyze transcriptional changes of *GhSWEET14a* and *GhSWEET14b* following inoculation of *Xcm* HM2.2 expressing individual TALEs from *Xcm* TX4. *Xcm* HM2.2 carrying Tal7b, but not other TALEs, significantly upregulated *GhSWEET14a,* comparable to the expression level induced by *Xcm* TX4 (**Figure 3A**). Similarly, *Xcm* HM2.2 carrying Tal7b, but not others, induced the expression of *GhSWEET14b* (**Figure 3B**). Importantly, *Xcm* HM2.2 carrying Tal7b, but not other TALEs, strongly induced water-soaked lesions (**Figure 3C and Supplementary Figure 3A**). These results indicate that Tal7b upregulates the expression of both *GhSWEET14a* and *GhSWEET14b* and mediates water-soaked lesion development.

**Figure 3.**
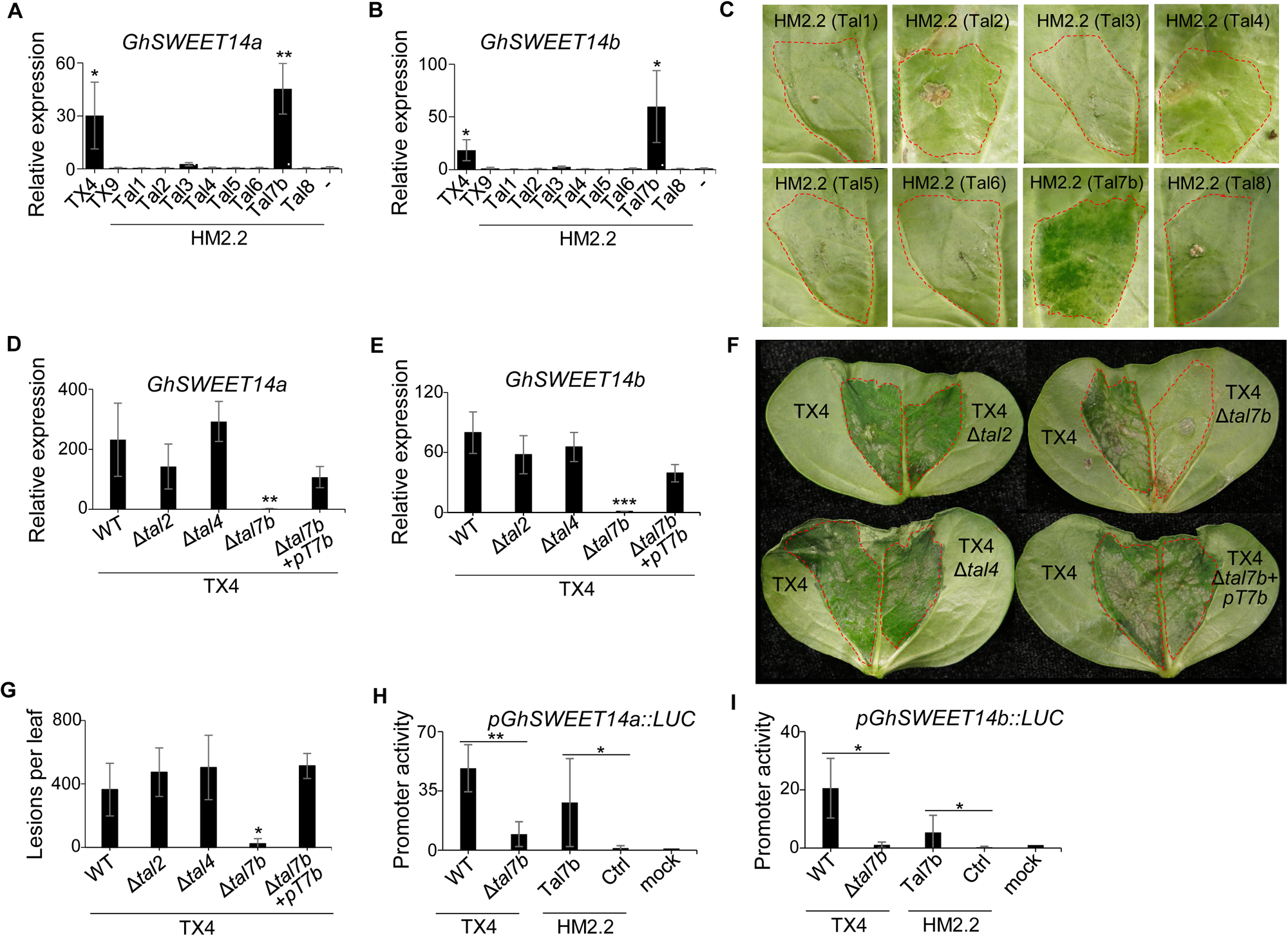
Tal7b induces the expression of *GhSWEET14a* and *GhSWEET14b* and mediates *Xcm* TX4-induced water-soaked symptoms. **(A & B)** Tal7b, but not other TALEs, in *Xcm* HM2.2, upregulates the expression of *GhSWEET14a* and *GhSWEET14b* in cotton. *Xcm* HM2.2, devoid of all TALEs, was transformed with a broad-host-range vector *pKEB1* carrying individual TALEs. *Xcm* TX4 and TX9 were included as controls. Bacterial suspensions of OD_600_ = 0.1 were syringe-inoculated into ten-day-old cotton cotyledons. RNA was extracted at 24 hours post-inoculation (hpi), and gene expression was analyzed by RT-qPCR. *GhUBQ* was used as an internal control. Data are shown as mean ± s.d. from three independent repeats (*n* = 3). Asterisks indicate a significant difference (* *p* < 0.05, ** *p* < 0.01)) compared to *Xcm* HM2.2 control (−) using a two-tailed Student’s *t*-test. **(C)** Tal7b, but not other TALEs, restores water-soaked symptom development to *Xcm* HM2.2. The bacterial inoculation was done as in (**A**). Photos were taken at 9 dpi. **(D & E)** *Xcm* TX4-induced *GhSWEET14a* and *GhSWEET14b* expression depends on Tal7b. *Xcm* TX4 deletion mutants, Δ*tal2*, Δ*tal4,* and *Δtal7b,* and *Δtal7b* carrying Tal7b in *pKEB1*, were syringe-inoculated into ten-day-old cotton cotyledons at OD_600_ = 0.1. RNA was extracted at 24 hpi, and gene expression was analyzed by RT-qPCR. *GhUBQ* was used as an internal control. Data are shown as mean ± s.d. (*n* = 3) from three independent repeats. Asterisks indicate a significant difference (** *p* < 0.01, *** *p* < 0.001) compared to wild-type (WT) *Xcm* TX4 control using a two-tailed Student’s *t*-test. **(F)** *Xcm* TX4-triggered water-soaking depends on Tal7b. The bacterial inoculation was done as in (**D**) with bacterial suspensions at OD_600_ = 0.05. Photos were taken at 6 dpi. **(G)** *Xcm* TX4-triggered water-soaked lesion number following vacuum infiltration depends on Tal7b. Bacterial suspensions at OD_600_ = 0.0001 were vacuum infiltrated into three-week-old cotton. Water-soaked lesions were counted on the 3^rd^ true leaf at 9 dpi. Data are shown as mean ± s.d. (*n* = 3) from three independent repeats. Asterisks indicate a significant difference (* *p* < 0.05) compared to WT *Xcm* TX4 control using a two-tailed Student’s *t*-test. **(H & I)** Tal7b activates *pGhSWEET14a::LUC* and *pGhSWEET14b::LUC* promoters in *N. benthamiana*. Leaves from five-week-old *N. benthamiana* were syringe-infiltrated with *Agrobacterium* carrying *pGhSWEET14a::LUC* (**H**) or *pGhSWEET14b::LUC* (**I**) at OD_600_ = 1.5. After 24 hours, suspensions of *Xcm* TX4, TX4Δ*tal7b*, *Xcm* HM2.2 carrying Tal7b or *pKEB1* empty vector (Ctrl.), or H_2_O mock controls were syringe-inoculated into the same area at OD_600_ = 0.5. Luciferase activity was quantified from co-inoculated leaf tissues harvested after another 24 hours and normalized to the H_2_O mock control (set as 1). Data are shown as mean ± s.d. (*n* = 4) from four independent repeats. Asterisks indicate a significant difference (* *p* < 0.05, ** *p* < 0.01) using a two-tailed Student’s *t*-test. The experiments were repeated three times with a similar result.

Next, we investigated whether *Xcm* mutants with Tal2, Tal4, or Tal7b missing are less virulent in cotton by generating a series of TX4 TALE mutant strains (**Supplementary Figures 3B and 3C**). RT-qPCR analysis showed that *Xcm* TX4Δ*tal7b*, but not TX4Δ*tal2* or TX4Δ*tal4*, induced *GhSWEET14a* and *GhSWEET14b* significantly less than wild-type (WT) TX4 (**Figures 3D and 3E**). Importantly, the capacity to induce *GhSWEET14a* and *GhSWEET14b* was restored upon introducing *pKEB1* carrying Tal7b into *Xcm* TX4Δ*tal7b* (**Figures 3D and 3E**). Furthermore, unlike *Xcm* TX4, TX4Δ*tal7b* could not induce the water-soaked phenotype in cotton when infiltrated into the leaf, and water-soaking was restored by complementation with *pKEB1-Tal7b* (**Figure 3F**). The deletion of *Tal2* or *Tal4*, TX4Δ*tal2* or TX4Δ*tal4*, did not affect water-soaked lesion development compared to *Xcm* TX4 (**Figure 3F**). The strains were also evaluated by counting the number of lesions that developed following vacuum-inoculation. The number of water-soaked lesions was significantly reduced upon inoculation with *Xcm* TX4Δ*tal7b* compared to TX4, TX4Δ*tal2*, and TX4Δ*tal4* (**Figure 3G**) and *pKEB1-Tal7b* fully complemented this phenotype (**Figure 3G**). Notably, *in planta* bacterial growth assay showed no significant difference between *Xcm* TX4 and TX4Δ*tal7b* populations over the course of seven days (**Supplementary Figure 3D**). Thus, Tal7b triggers the elevated expression of *GhSWEET14a* and *GhSWEET14b* and elicits water-soaked lesions but does not affect bacterial multiplication *in planta*. This resembles the function of Avrb6 in *Xcm* H1005, which enhances water-soaked lesion development by inducing *GhSWEET10* expression but does not contribute to bacterial growth in cotton leaves^21,28^.

The induction of *GhSWEET14a* and *GhSWEET14b* by Tal7b prompted us to test whether their promoters, fused with the firefly luciferase (*LUC*) reporter gene, could serve as a sensitive reporter for monitoring *Xcm* infections. Individually, we fused the 1-kb regions upstream of the translational start site of *GhSWEET14a* and *GhSWEET14b* with *LUC* and assayed these constructs by *Agrobacterium*-mediated transient transformation in *Nicotiana benthamiana* followed 24 hr later by infiltration of *Xcm* TX4 or HM2.2 carrying Tal7b. Each *Xanthomonas* strain significantly activated both *pGhSWEET14a::LUC* and *pGhSWEET14b::LUC* compared to mock treatment or *Xcm* HM2.2 alone (**Figures 3H and 3I**). The *Xcm* TX4Δ*tal7b* inoculation resulted in a significantly lower induction of *pGhSWEET14a::LUC* and *pGhSWEET14b::LUC* promoters than WT TX4 (**Figures 3H and 3I**). Together, our comprehensive gain-of-function and loss-of-function assays of individual TALEs indicate that Tal7b from *Xcm* TX4 plays a vital role in the induction of *GhSWEET14a* and *GhSWEET14b* genes, as well as water-soaked lesion development in cotton.

### *GhSWEET14a*/*b* mediate Tal7b-triggered disease susceptibility

To determine the role of *GhSWEET14a* and *GhSWEET14b* in Tal7b-triggered water-soaked lesion development, we silenced *GhSWEET14a* and *GhSWEET14b* in cotton by *Agrobacterium*-mediated virus-induced gene silencing (VIGS) followed by inoculations of *Xcm* TX4 or HM2.2 carrying Tal7b (**Figure 4A**). The tetraploid upland cotton contains A and D subgenomes^37^. *GhSWEET14a* has two homoeologous genes in both A and D subgenomes, referred to as *GhSWEET14aA* and *GhSWEET14aD,* respectively, and they share 99.1% sequence identity at the nucleotide level (**Supplemental Figure 4A**). *GhSWEET14b* is only present in the D subgenome, and thus referred to as *GhSWEET14bD* (**Supplemental Figure 4A**)*. GhSWEET14bD* is 80.5% and 80.6% identical at the nucleotide level to *GhSWEET14aA* and *GhSWEET14aD*, respectively. We constructed a VIGS vector, VIGS-*GhSWEET14a/b*, that concurrently reduced the expression of *GhSWEET14a* (including both *GhSWEET14aA* and *GhSWEET14aD*) and *GhSWEET14b,* in comparison to the vector control (**Supplemental Figure 4B**). Once gene silencing was established, *Xcm* TX4 or HM2.2 carrying Tal7b was syringe-inoculated into cotton leaves. Compared to those inoculated with the VIGS vector control, the *GhSWEET14a/b-*silenced cotton leaves exhibited a noticeable reduction in water-soaked symptoms caused by *Xcm* TX4, confirming the important role of *GhSWEET14a/b* in water-soaked symptom development (**Figure 4A, top panel**). Furthermore, the water-soaked symptom development triggered by *Xcm* HM2.2 carrying Tal7b was also reduced in the *GhSWEET14a/b-*silenced cotton leaves compared to VIGS vector-control leaves (**Figure 4A, bottom panel**). Moreover, *GhSWEET14a/b*-silenced cotton leaves vacuum-infiltrated with *Xcm* TX4 or HM2.2 carrying Tal7b showed fewer water-soaked lesions than VIGS control leaves inoculated with those strains (**Figures 4B and 4C**).

**Figure 4.**
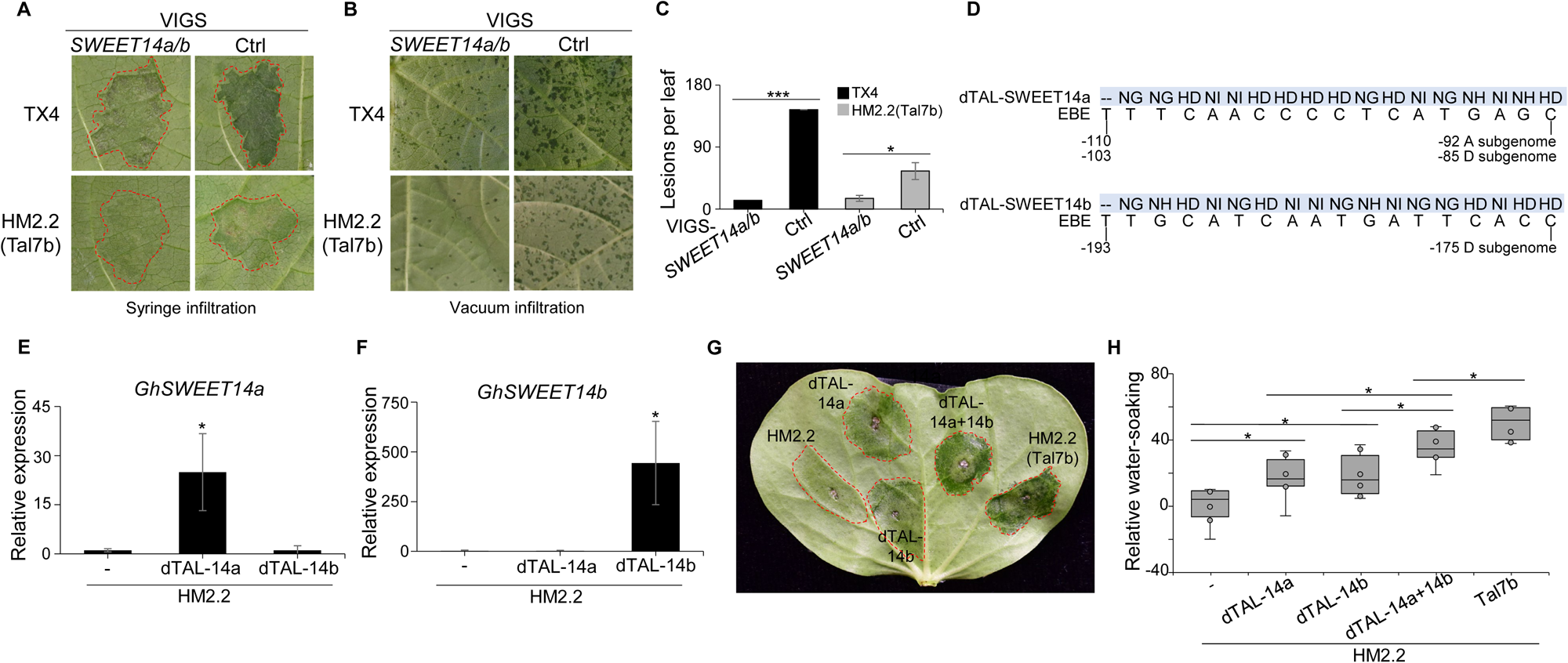
*GhSWEET14a* and *GhSWEET14b* collectively contribute to *Xcm*-induced water-soaked lesions in cotton. **(A)** Silencing *GhSWEET14a* and *GhSWEET14b* reduces the severity of *Xcm* TX4 and HM2.2 Tal7b-induced water-soaked symptoms. Seven-day-old cotton cotyledons were syringe-inoculated with *Agrobacterium* carrying VIGS constructs to silence both *GhSWEET14a* and *GhSWEET14b*, or a vector control. Three weeks later, when silencing was established (**Supplementary Figure S4**), the 2^nd^ true leaves were syringe-inoculated with *Xcm* TX4 or HM2.2 carrying Tal7b at OD_600_ = 0.1. Photos were taken at 5 dpi. **(B & C)** Silencing *GhSWEET14a* and *GhSWEET14b* reduces the number of lesions following vacuum infiltration of *Xcm* TX4 and HM2.2 Tal7b. Bacterial suspensions at OD_600_ = 0.0001 were vacuum infiltrated into true leaves of three-week-old cotton plants. Photos were taken (**B**), and water-soaked lesions were counted (**C**) on the 2nd true leaves at 10 dpi. Data are shown as mean ± s.e. (*n* = 5) from five independent repeats. Asterisks indicate a significant difference (* *p* < 0.05, *** *p* < 0.001) in *GhSWEET14a*/*b*-silenced plants compared to vector controls using a two-tailed Student’s *t*-test. **(D)** RVD sequences of designer TALEs (dTALE) and their corresponding predicted EBE sites within the promoters of *GhSWEET14a* and *GhSWEET14b*. The numbers below indicate the EBE locations in proximity to the translational start codon of *GhSWEET14a* or *GhSWEET14b*. dTAL-SWEET14a targets an EBE in the promoter of the gene in both the A and D subgenomes, distinct from the Tal7b EBE, and dTAL-SWEET14b targets an EBE in the promoter of the single copy of the gene, present in the D subgenome. **(E & F)** The dTALEs can effectively and specifically induce the expression of target genes. Bacterial suspensions of *Xcm* HM2.2 carrying dTAL-SWEET14a, dTAL-SWEET14b, or vector control were syringe-inoculated into seven-day-old cotton cotyledons at OD_600_ = 0.1. RNA was extracted at 24 hpi, and gene expression was analyzed by RT-qPCR. *GhUBQ* was used as an internal control. Data are shown as mean ± s.d. (*n* = 3) from three independent repeats. Asterisks indicate a significant difference (* *p* < 0.05) compared to vector control using a two-tailed Student’s *t*-test. **(G & H)** Upregulation of *GhSWEET14a* and *GhSWEET14b* by dTALEs induces water-soaked symptoms, in an incompletely additive, partly redundant manner. Bacterial suspensions of *Xcm* HM2.2 carrying dTAL-SWEET14a, dTAL-SWEET14b, or both were syringe-inoculated into seven-day-old cotton cotyledons at OD_600_ = 0.1. Photos were taken at 5 dpi (**G**). Water-soaked lesions were analyzed using ImageJ FIJI (**H**). Data are shown as the negative grayscale mean of water-soaked lesions compared to *Xcm* HM2.2 vector control. The gray-scale mean was determined by averaging the means of gray-scale values in the RGB channels from 6 independent repeats. Asterisks indicate a significant difference (* *p* < 0.05) between treatments using a two-tailed Student’s *t*-test. The experiments were repeated three times with a similar result.

Gene-specific dTALEs were engineered to individually induce the expression of *GhSWEET14a* or *GhSWEET14b.* The best DNA sequence match to the RVD sequence of Tal7b in each gene was predicted using the Target Finder tool of TALE-NT 2.0^38^, and the dTALEs were designed to target sequences distinct from that candidate EBE in each gene promoter (**Figure 4D**). RT-qPCR confirmed the specific induction of *GhSWEET14a* by dTAL-SWEET14a and *GhSWEET14b* by dTAL-SWEET14b when expressed in *Xcm* HM2.2 (**Figures 4E and 4F**). Notably, HM2.2 harboring either dTALE alone caused a water-soaked phenotype, but to a lesser extent than HM2.2 carrying Tal7b (**Figures 4G and 4H**). Importantly, co-inoculating *Xcm* HM2.2 carrying dTAL-SWEET14a and dTAL-SWEET14b induced a significantly stronger water-soaked phenotype than the strains individually, yet still at a lower level than Tal7b and less than completely additively, suggesting partially redundant function with respect to susceptibility (**Figures 4G and 4H**). The data indicate that *GhSWEET14a* and *GhSWEET14b* contribute quantitatively and partially redundantly to the development of water-soaked symptoms, but they also hint at one or more additional *S* gene(s) targeted by Tal7b.

### *GhSWEET14a*/*b* play dual roles in *Xcm* virulence and cotton boll development

To further investigate the roles of *GhSWEET14a* and *GhSWEET14b*, we targeted *GhSWEET14a* and *GhSWEET14b* for mutagenesis through multiplexed CRISPR-Cas9-mediated editing. The gRNA-*GhSWEET14a/b* construct was designed to simultaneously target *GhSWEET14a* and *GhSWEET14b* in both the A and D subgenomes. Given the sequence conservation within upland cotton A and D subgenomes, gRNA-*GhSWEET14a* targeted both *GhSWEET14aA* and *GhSWEET14aD*. As *GhSWEET14b* is only present in the D subgenome, gRNA-*GhSWEET14b* targeted *GhSWEET14bD*. These gRNAs were arranged in a tandem array of tRNA-gRNA motifs expressed as a single polycistronic tRNA-gRNA gene^39^. Cleavage of primary transcripts by the endogenous tRNA-processing system releases mature gRNA molecules^40^.

gRNA-*GhSWEET14a/b*, along with the *Cas9* gene in the *pRGEB32-GhU6.9* vector^39^, was transformed into the cotton variety Coker 312 through standard tissue culture regeneration to generate *GhSWEET14a/b*-edited lines. Among five lines we screened by targeted sequencing of *GhSWEET14a/b*, line 5-2 contained a 2-nucleotide (nt) deletion in the first exon of both *GhSWEET14aA* and *GhSWEET14aD,* as well as a 1-nt deletion in the first exon of *GhSWEET14bD*, each resulting in a functionally disruptive frameshift (**Figure 5A**). Significantly, following *Xcm* TX4 inoculation, water-soaked lesion development was markedly reduced in the CRISPR-Cas9-*GhSWEET14a/b* edited cotton line compared to the control line transformed with GFP (**Figure 5B**). The result substantiates the crucial role of *GhSWEET14a* and *GhSWEET14b* in *Xcm* TX4 virulence and suggests a promising strategy for conferring bacterial blight resistance in allotetraploid cotton by simultaneously targeting multiple *S* genes.

**Figure 5.**
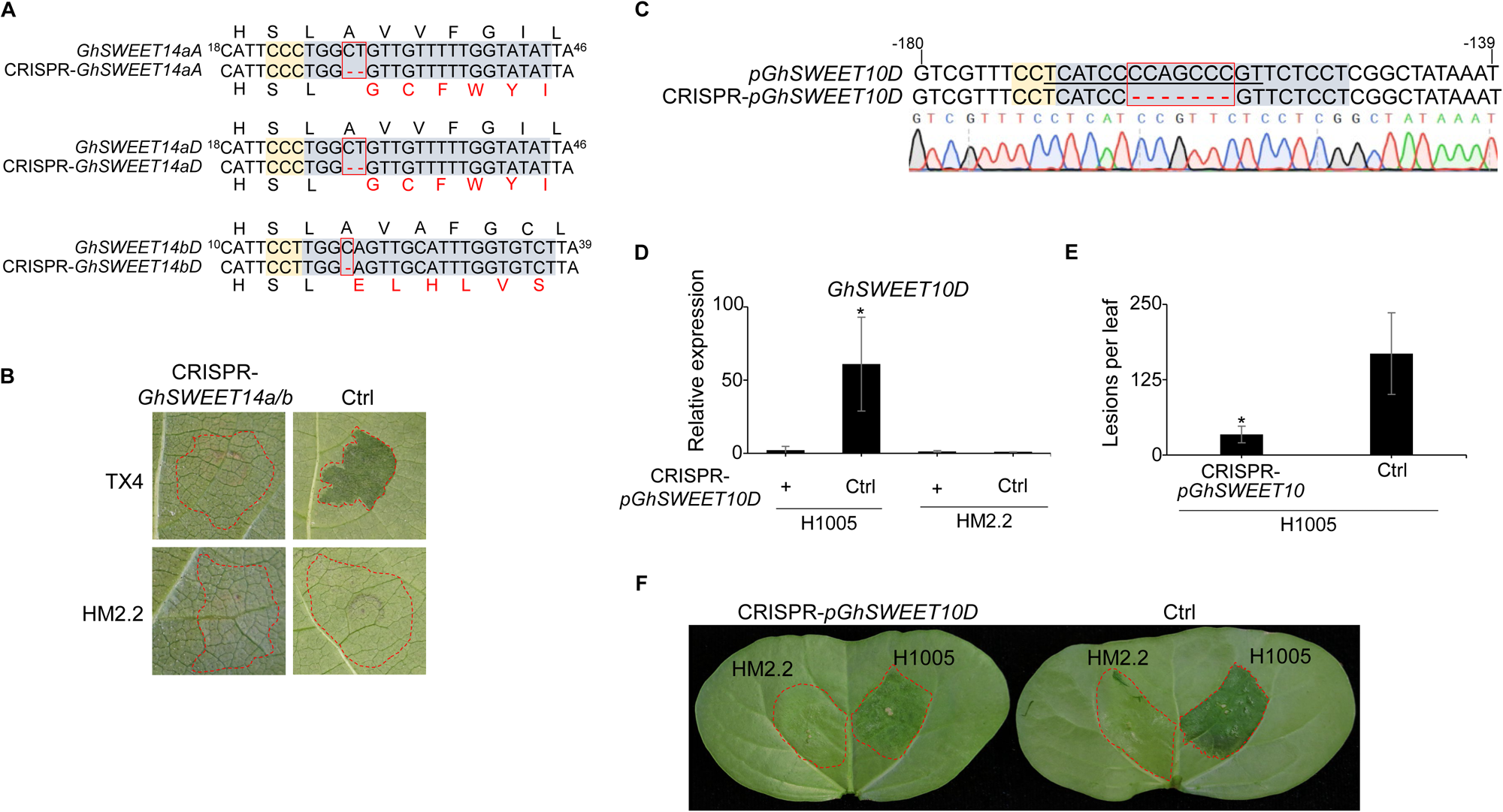
CRISPR-Cas9-mediated gene editing of *GhSWEET* genes reduces *Xcm*-induced water-soaked lesion development. **(A)** The CRISPR-*GhSWEET14a/b* gene edited cotton line harbors early frameshift deletions in both *GhSWEET14a* and *GhSWEET14b*. Genomic DNA was extracted from the edited line, and the target site was amplified by PCR and the amplicon sequenced. Nucleotide deletions are denoted by red dashes. Amino acid sequences are displayed above and below, with frameshift mutations indicated by red letters. Shaded boxes highlight gRNA target sequences and PAM sites. Numbers on the sides indicate the target site’s proximity to the translational start codon. **(B)** Water-soaked symptom development is reduced in CRISPR-*GhSWEET14a/b* transgenic cotton compared to vector control transgenic cotton upon *Xcm* TX4 infection. Transgenic cotton plants were syringe-inoculated with bacterial solutions of *Xcm* TX4 or HM2.2 at OD_600_ = 0.05. Photos were taken at 5 dpi. **(C)** The CRISPR-*pGhSWEET10D* line contains a 7-nucleotide deletion in the *GhSWEET10D* promoter at the Avrb6 EBE, determined by PCR amplification and sequencing. Nucleotide deletions are denoted by red dashes. A sequencing chromatogram of the targeted region is shown below the nucleotides. Shaded boxes highlight the gRNA target sequence and PAM site. The underlined sequence is the Avrb6 EBE. Numbers on the sides indicate the target site’s proximity to the translational start codon. **(D)** *Xcm* H1005-mediated upregulation of *GhSWEET10D* is reduced in the CRISPR-*pGhSWEET10D* line. Ten-day-old transgenic cotton cotyledons were syringe-inoculated with bacterial solutions of *Xcm* H1005 or HM2.2 at OD_600_ = 0.1. RNA was extracted at 24 hpi, and gene expression was analyzed by RT-qPCR. *GhUBQ* was used as an internal control. Data are shown as mean ± s.d. (*n* = 3) from three independent repeats. Asterisks indicate a significant difference (* *p* < 0.05) compared to the control line using a two-tailed Student’s *t*-test. **(E)** Lesion development induced by *Xcm* H1005 is reduced in the CRISPR-*pGhSWEET10D* line. Twenty-one-day-old transgenic cotton plants were vacuum-infiltrated with *Xcm* H1005 at OD_600_ = 0.0001. Water-soaked lesions were counted on the 3^rd^ true leaf at 10 dpi. Data are shown as mean ± s.e. (*n* = 6) from six independent repeats. Asterisks indicate a significant difference (* *p* < 0.05) compared to the control line using a two-tailed Student’s *t*-test. **(F)** The CRISPR-*pGhSWEET10D* line exhibits reduced water-soaked symptoms following *Xcm* H1005 infection. Ten-day-old transgenic cotton cotyledons were syringe-inoculated with bacterial solutions of *Xcm* H1005 or HM2.2 at OD_600_ = 0.1. Photos were taken at 4 dpi. The experiments were repeated three times with a similar result.

However, it’s noteworthy that the CRISPR-*GhSWEET14a/b* plants did not produce bolls. We observed a striking contrast in reproductive development between the CRISPR-*GhSWEET14a/b* plants and control plants. While the control plants exhibited typical flower withering and subsequent boll formation a few days after bloom, the flowers of the CRISPR-*GhSWEET14a/b* plants persisted for over 20 days without any observable boll development (**Supplementary Figure 5A**) This phenomenon corroborates the significance of sugar transporters, including *SWEET* genes, in various development and reproduction stages of plants^13^. The absence of boll development in the CRISPR-*GhSWEET14a/b* plants suggests the essential nature of *GhSWEET14a* and *GhSWEET14b* in cotton development and reproduction.

### Gene editing of the *GhSWEET10* promoter EBE for Avrb6 confers resistance to *Xcm* H1005

As illustrated above and observed by others, knockout of *SWEET* genes can be detrimental (**Supplementary Figure 5A**)^25,41^. Mutagenesis of TALE EBE sites in the *S* genes could offer a strategy for achieving resistance without compromising growth or reproduction^42,43^. As a proof-of-concept in cotton, we designed a gRNA-*pGhSWEET10* construct to target the EBE of *GhSWEET10,* the *S* gene induced by Avrb6 in *Xcm* H1005^21^. Notably, the induction of *GhSWEET10D* by Avrb6 is much higher than its A subgenome homoeolog *GhSWEET10A,* indicating *GhSWEET10D* as a primary *S* gene in *Xcm* H1005 infection^21^.

Through CRISPR-Cas9-mediated gene editing, we successfully introduced a 7-nucleotide deletion in the Avrb6 EBE within the *GhSWEET10D* promoter in a cotton line designated CRISPR-*pGhSWEET10D* (**Figure 5C**). RT-PCR analysis revealed that the CRISPR-*pGhSWEET10D* line maintained a comparable basal level of *GhSWEET10D* transcripts to the cotton line transformed with GFP as controls (**Supplementary Figure 5B**), indicating undisturbed endogenous expression of *GhSWEET10D* in the CRISPR-*pGhSWEET10D* line. Importantly, *Xcm* H1005-mediated induction of *GhSWEET10D* was almost completely abrogated in the CRISPR-*pGhSWEET10D* line compared to the control line (**Figure 5D**). The induction of *GhSWEET10D* by *Xcm* H1005 in the CRISPR-*pGhSWEET10D* line was similar to that by *Xcm* HM2.2, which lacks Avrb6 (**Figure 5D**), suggesting that Avrb6 delivered by *Xcm* H1005 was ineffective at inducing *GhSWEET10D* in this line.

Furthermore, the CRISPR-*pGhSWEET10D* line displayed a significant reduction in water-soaked lesion development and lesion count following either hand-or vacuum-inoculation with *Xcm* H1005, in comparison to the control line (**Figures 5E and 5F**). Notably, bacterial counting assays following *Xcm* H1005 infection showed no significant difference in bacterial populations between the CRISPR-*pGhSWEET10D* line and the control line over a seven-day period (**Supplementary Figure 5C**). This lack of effect on bacterial multiplication aligns with our prior results obtained by knockdown of *GhSWEET10* using VIGS^21^. Importantly, the CRISPR-*pGhSWEET10D* line displayed no growth defects, with normal boll development. These results highlight the potential of CRISPR-Cas9-mediated disruption of *S* gene EBEs rather than coding sequences as a strategy for controlling BBC.

### Integration of transcriptome analysis and EBE site prediction reveals *GhSWEET14a/b* as Tal7b targets

To uncover possible additional targets of Tal7b, we performed whole-genome RNA-sequencing (RNA-seq) analysis using the Illumina HiSeq 2000 platform to identify Tal7b-induced *G. hirsutum* genes. Cotyledons from the BBC susceptible cotton variety FiberMax FM 2322GL (FM 2322GL) were inoculated with either *Xcm* HM2.2 carrying Tal7b or an empty vector before being harvested 24 hours post-inoculation (hpi). Our analysis identified 707 genes induced by *Xcm* HM2.2 Tal7b compared to the *Xcm* HM2.2 control using cutoffs of log_2_ fold change > 1 and *p*-value < 0.05 (**Figure 6A**). Next, we screened the *G. hirsutum* promoterome for potential Tal7b EBEs using the TALE-NT 2.0 Target Finder tool^38^ and further manually checked these candidates with knowledge of the codes, including the context of the EBE in the promoter. Since the score ratios for the Tal7b candidate EBEs in the *GhSWEET14a/b* promoters are greater than 3.0 (**Figure 6B**), typically considered indicative of a weak candidate EBE, we relaxed the score ratio cutoff in this screen to 4.0. This analysis identified 397 genes containing predicted Tal7b EBEs within their promoters (**Figure 6A)**. By cross-referencing 707 Tal7b-induced genes with the 397 genes containing predicted Tal7b EBEs within their promoters, we uncovered 26 candidate gene targets of Tal7b (**Figure 6A, Supplementary Figure 7**). Among the top ten candidates, *GhSWEET14aA* and *GhSWEET14bD* exhibit the EBE score ratios with EBE positions around 100 bp upstream of the start codon and were induced approximately 20-fold by *Xcm* HM2.2 Tal7b compared to *Xcm* HM2.2 (**Figures 6B and 6C**). Given the molecular and physiological evidence we obtained for *GhSWEET14a* and *GhSWEET14b* as targets of Tal7b, these findings underscore the feasibility of RNA-seq analysis coupled with EBE site prediction in identifying TALE target genes. However, they also emphasize the fact that EBE score ratios for valid targets may exceed the commonly used cutoff for the prediction of candidates. In addition to the Tal7b EBEs in the *GhSWEET14a/b* gene promoters described here, this is the case for the EBE of the *X. oryzae* pv. *oryzae* TALE AvrXa27 in the promoter of the rice *Xa27* gene^44^.

**Figure 6.**
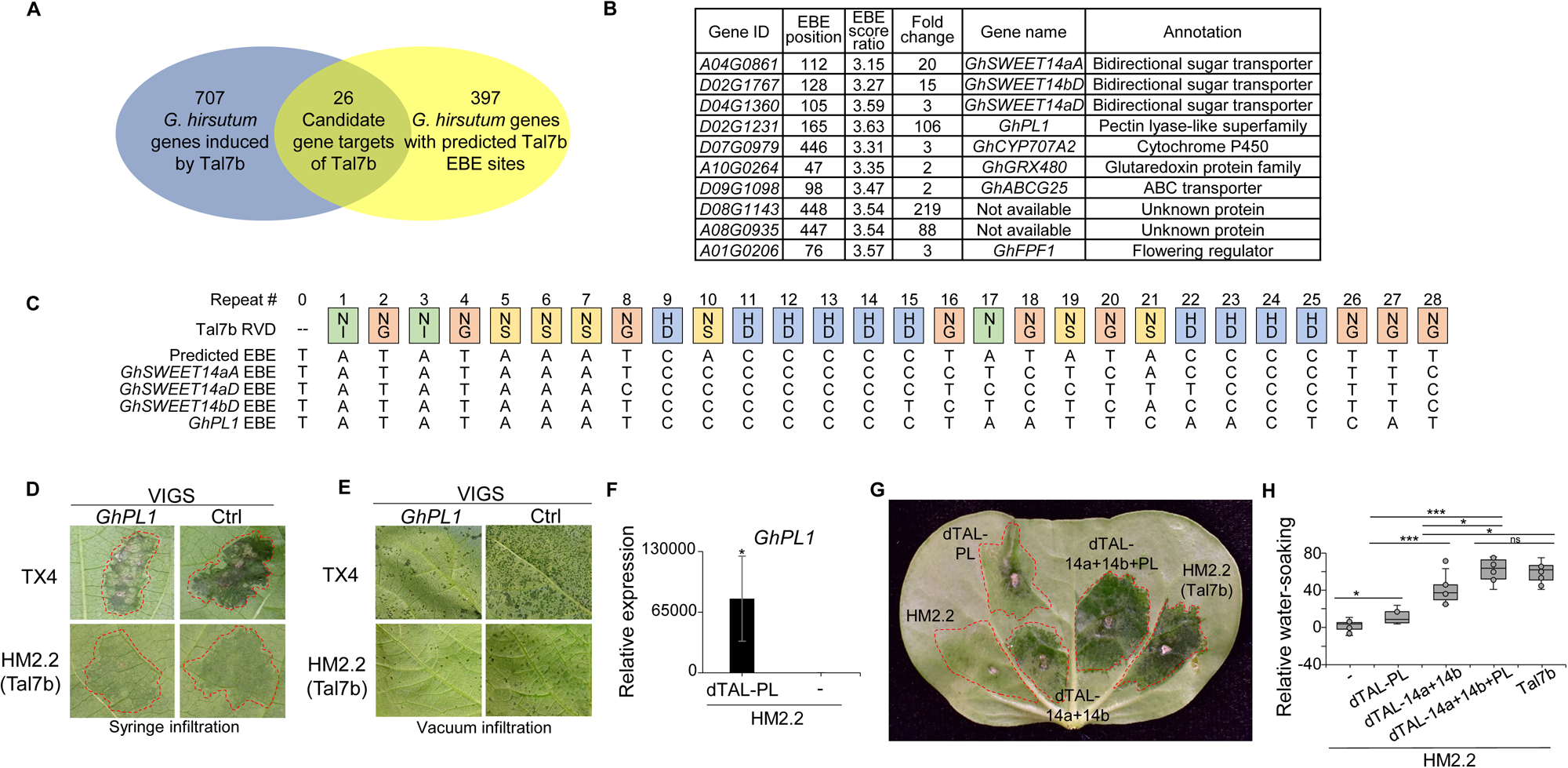
Transcriptome profiling coupled with EBE prediction reveals a pectin lyase as an additional target of Tal7b, contributing to *Xcm* water-soaked lesion development. **(A)** Venn diagram of Tal7b-induced genes by RNA-sequencing (RNA-seq) analysis and Tal7b-candidate target genes by *in silico* EBE prediction using TALE-NT 2.0 with a relaxed score ratio cutoff, 4.0. **(B)** The top ten candidate genes induced by Tal7b ranked by the strength of the EBE score ratio, the promoter context of the EBE^38^ and induction fold. EBE position indicates its proximity to the translational start codon. **(C)** The Tal7b RVD sequence and its predicted EBE sites. Colored boxes depict the RVDs of each repeat motif of Tal7b, with the number above each box indicating the RVD position within the CRR. The Tal7b EBE sequences predicted in the promoters of *GhSWEET14a*, *GhSWEET14b*, and *GhPL1* using TALE-NT 2.0^38^ are listed below the RVDs. **(D)** Silencing *GhPL1* reduces water-soaked symptoms caused by *Xcm* TX4 or HM2.2 carrying Tal7b. Seven-day-old cotton cotyledons were syringe-inoculated with *Agrobacterium* carrying VIGS constructs to silence *GhPL1* or with *Agrobacterium* carrying a vector control. Three weeks later, when silencing was established, the 2^nd^ true leaves were syringe-inoculated with *Xcm* TX4 or HM2.2 Tal7b at OD_600_ = 0.1. Photos were taken at 5 dpi. **(E)** Silencing *GhPL1* reduces the water-soaked lesion development caused by *Xcm* TX4 or HM2.2 carrying Tal7b. Bacterial suspensions at OD_600_ = 0.0001 were vacuum infiltrated into three-week-old cotton. Photos were taken on the 2^nd^ true leaves at 9 dpi. **(F)** *GhPL1* is significantly induced by dTALE dTAL-PL. Bacterial suspensions of *Xcm* HM2.2 carrying dTAL-PL or a vector control at OD_600_ = 0.1 were syringe-inoculated into seven-day-old cotton cotyledons. RNA was extracted at 24 hpi, and gene expression was analyzed by RT-qPCR. *GhUBQ* was used as an internal control. Data are shown as mean ± s.d. (*n* = 3) from three independent repeats. Asterisks indicate a significant difference (* *p* < 0.05) compared to the vector control using a two-tailed Student’s *t*-test. **(G & H)** *GhPL1* quantitatively contributes to the water-soaked symptom development, synergistically with *GhSWEET14a/b.* Bacterial suspensions or co-suspensions of *Xcm* HM2.2 carrying dTAL-PL, dTAL-SWEET14a and dTAL-SWEET14b, or all three dTALEs were syringe-inoculated into seven-day-old cotton cotyledons at OD_600_ = 0.1. Equal proportions of each bacterial inoculum were combined with an empty vector co-inoculum. *Xcm* HM2.2 carrying Tal7b, co-inoculated with HM2.2 carrying empty vector to maintain the same relative OD_600_ as each of the others, was included as a control. Photos were taken (**H**), and water-soaked lesions were analyzed using ImageJ FIJI (**I**) at four dpi. Asterisks indicate a significant difference (* *p* < 0.05, *** *p* < 0.001, ns = no significance) between treatments using a two-tailed Student’s *t*-test. The experiments were repeated three times with a similar result.

### A pectin lyase functions synergistically with *GhSWEET14a/b* to induce water-soaked lesions

Three genes in the top 10 were more strongly induced by Tal7b than *GhSWEET14aA* (*A04G0861*) and *GhSWEET14bD* (*D02G1767*), as much as ∼200-fold. Two of these, *GhD08G1143* and *GhA08G0935*, encode proteins of unknown function. Notably, their predicted EBE sites are positioned more than 400 bp upstream from the start codon, a location typically deemed less likely as a target gene^45^. The third, *GhD02G1231*, with an over 100-fold change in expression, encodes a pectin lyase-like protein, which was termed *GhPL1*. To investigate whether the expression of *GhPL1* contributes to BBC disease symptom development, we silenced it with *Agrobacterium*-mediated VIGS, followed by *Xcm* inoculations. Notably, following syringe-inoculation with *Xcm* TX4 or HM2.2 carrying Tal7b, *GhPL1*-silenced cotton exhibited reduced water-soaked symptoms compared to a non-silenced vector-only control (**Figure 6D**). Similarly, *GhPL1*-silenced cotton produced significantly fewer water-soaked lesions than the vector-only control when vacuum-infiltrated with *Xcm* TX4 or HM2.2 carrying Tal7b (**Figure 6E**). These results suggest that the induction of *GhPL1* also contributes to the development of water-soaked lesions.

To further test this conclusion, a dTALE, dTAL-PL, was constructed to induce the expression of *GhPL1* through an EBE distinct from that of Tal7b. RT-qPCR analysis confirmed the effective induction of *GhPL1* by HM2.2 expressing dTAL-PL (**Figure 6F**). The upregulation of *GhPL1* by dTAL-PL was associated with weak water-soaking, intermediate to the barely discernible water-soaking induced by HM2.2 with an empty vector and the more obvious water-soaking induced by co-inoculated transformants of HM2.2 expressing dTAL-SWEET14a and dTAL-SWEET14b (**Figures 6G and 6H**). To assess the relationship of *GhPL1* and *GhSWEET14a*/*b* in the development of BBC disease symptoms, we compared the severity of the water-soaking caused by dTAL-PL, dTAL-SWEET14a and dTAL-SWEET14b together, and all three in combination, relative to that caused by Tal7b. Importantly, visually and when quantified by image analysis using ImageJ, water-soaking was strongest when all three dTALEs were tested in combination, reaching a level comparable to that caused by Tal7b (**Figures 6G and 6H**). Further, the water-soaking caused by all three dTALEs in combination was greater than the sum of the water-soaking caused by dTAL-PL alone and dTAL-SWEET14a and dTAL-SWEET14b together. Thus, it is likely that the pectin lyase plays an integral role in the development of BBC disease symptoms synergistically with *GhSWEET14a* and *GhSWEET14b*.

## DISCUSSION

Identifying resistance (*R*) or susceptibility (*S*) genes in tetraploid cotton remains a challenge. Initially, *GhSWEET10* was identified as an *S* gene targeted by Avrb6 in the early *Xcm* H1005 isolate from BBC-infected cotton fields, but its current relevance to protection against BBC is uncertain, as *GhSWEET14a* and *GhSWEET14b* rather than *GhSWEET10* are induced by reemerging *Xcm* isolates^21^. In this study, we identified Tal7b in a collection of *Xcm* isolates from Texas, USA, targeting *GhSWEET14a* and *GhSWEET14b* to promote disease, specifically water-soaked symptoms. Despite the overall similarity in genome sequence and organization between *Xcm* H1005 and reemerging Texas isolates, their TALomes exhibit distinct RVD compositions, suggesting rapid evolution of TALEs associated with virulence shifts over time. However, the TALomes of three Texas *Xcm* isolates, TX4, TX7, and TX9, exhibit a high degree of overall sequence identity and share similar genome/plasmid locations among them, as well as with those of an isolate from Mississippi, MSCT1. Although the whole genome sequence is unavailable, the TALEs in *Xcm* Xss-V2-18, a highly virulent strain in China, closely resemble those in *Xcm* TX4 and MSCT1^46^. Among six TALEs in *Xcm* Xss-V2-18, four of them, Tal4, Tal1, Tal5, and Tal3, share identical RVD sequences with Tal3, Tal5, Tal6, and Tal8, respectively, in *Xcm* TX4^46^. Notably, *Xcm* Xss-V2-18 lacks Tal7b, crucial for *Xcm* TX4 virulence. Conversely, Tal2 in *Xcm* Xss-V2-18, responsible for its virulence, exhibits two mismatches compared to the corresponding Tal2 in *Xcm* TX4. This suggests that these reemerging *Xcm* isolates likely originated from a recent common ancestral but have developed distinct virulence strategies to infect cotton in different regions.

*SWEET* genes are common susceptibility gene targets of TALEs across several *Xanthomonas* spp. Multiple TALEs from different *X. oryzae* pv. *oryzae* strains, including AvrXa7, PthXo3, TalC, and Tal5, target *OsSWEET14* to promote infection^19,26^. In some cases, different TALEs within the same *X. oryzae* pv. *oryzae* strain target functionally redundant *SWEET* genes, as seen with AvrXa7 and PthXo2 from strain MAFF311018 targeting *OsSWEET14* and *OsSWEET13*, respectively^16,47^. In cassava bacterial blight, Tal20 of *X. axonopodis* pv. *manihotis* promotes disease by activating *MeSWEET10a*^48^. And, we previously demonstrated in cotton bacterial blight that *Xcm* strain H1005 depends on Avrb6 targeting *GhSWEET10D* to cause water-soaking^21^. In the present study, we demonstrate that Tal7b found in *Xcm* strain TX4 and other reemerging isolates simultaneously induces *GhSWEET14a* and *GhSWEET14b,* leading to the partially additive promotion of water-soaked lesions in cotton. This observation illustrates a new model relative to the ones mentioned above, in which a single effector induces the expression of multiple *SWEET* genes to achieve a high level of *SWEET* expression, crucial for SWEET-mediated susceptibility. The targeting and induction of both *GhSWEET14a* and *GhSWEET14b* in cotton by Tal7b suggest an adaptive mechanism whereby the pathogen efficiently guards against host evolution of a TALE-insensitive *S* gene allele by simultaneously targeting two functionally similar *S* genes with the same TALE.

It has been suggested that TALE-mediated sucrose efflux in the apoplast serves as a carbon source for bacteria, aiding in their survival and proliferation^16,19,20,23–26^. However, the Tal7b-mediated induction of *GhSWEET14a* and *GhSWEET14b* does not appear to influence bacterial multiplication *in planta*. Similar results were observed with Avrb6-mediated induction of *GhSWEET10D*, which does not affect *in planta* multiplication of *Xcm* H1005^21,28^. One explanation for this observation is that sucrose efflux might affect the osmotic potential of the apoplast, contributing to water-soaked lesions and benefiting bacterial virulence by regulating stomatal movement. Stomatal pores serve as entry points for many pathogens to invade leaf apoplasts^49^. Once inside, the aqueous environment within the leaf apoplast promotes pathogen colonization^50,51^. Furthermore, a high level of sucrose could induce stomatal closure^52^, and thereby increase water potential in the apoplast. Notably, the TALE AvrHah1 from *X. gardneri* also contributes to water-soaked lesion development but does not affect bacterial multiplication *in planta*^53^. It was proposed that in activating bHLH transcription factor genes that in turn activate a pectate lyase, AvrHah1 modifies the plant cell wall to promote water uptake that facilitates bacterial egress from the apoplast to the leaf surface^53^. Finally, Tal2g of the rice bacterial leaf streak pathogen *X. oryzae* pv. *oryzicola* contributes to lesion expansion and bacterial egress to the leaf surface, without impacting total bacterial populations, by activating *OsSULTR3;6*, encoding a putative sulfate transporter^54,55^. These findings collectively suggest that TALEs may contribute to pathogen virulence by modulating apoplast water dynamics and osmotic potential to create a favorable environment for colonization and dissemination.

The integration of transcriptome analysis and EBE site prediction^54^ has become a widely utilized strategy for identifying *S* genes targeted by *Xanthomonas* spp. TALEs^21,56–60^. Through this approach, we identified the pectin lyase gene *GhPL1* as an additional target of Tal7b, quantitively contributing to water-soaked symptom development, synergistically with *GhSWEET14a* and *GhSWEET14b*. Pectin is a major component of the plant cell wall. Pectolytic enzymes in plants function in diverse processes, from cell wall remodeling to fruit ripening, and others. Degradation of pectin by secreted enzymes of soft rot bacteria in the genera *Pectobacterium* and *Dickeya* leads to plant tissue maceration, electrolyte loss, and cell death^61^. How *GhPL1* contributes to water-soaking is not clear, but as illustrated by AvrHah1 of *X. gardneri*, discussed above, TALE-mediated induction of pectolytic enzymes may be a virulence strategy common to several *Xanthomonas* species. In addition to the pectate lyase gene induced by *X. gardneri*, a TALE-targeted pectate lyase gene was identified as a candidate *S* gene in rice bacterial blight of rice^58^. Also, *X. axonopodis* pv. *manihotis* deploys a TALE, Tal14, that induces two pectate lyases^62^. Interestingly, however, dTALE experiments in that study suggested that this induction might actually be counter-adaptive, because the dTALEs resulted in restricted pathogen growth^59^. Our findings, validated through gain-of-function (dTALEs) and loss-of-function (VIGS) approaches, demonstrate that *GhPL1* functions as a *bona fide S* gene in cotton, contributing to water-soaked lesion development.

Our study adds to the growing body of literature showing the effectiveness of CRISPR-Cas9-mediated gene editing of *S* genes for resistance to diseases caused by *Xanthomonas* spp. Despite the complexities posed by the tetraploid genome of *G. hirsutum*, we successfully targeted *GhSWEET14a* in both subgenomes and the sole copy of *GhSWEET14b*, resulting in simultaneous knockouts of all three and consequent reduction of *Xcm*-mediated water-soaked symptoms. However, it is important to note that the edits also resulted in a defect in reproductive development, the failure to produce bolls. This observation aligns with findings in rice, where knockout of *OsSWEET11* led to defective grain filling and abnormal pollen development^41^, underscoring the critical roles of at least some *SWEET* genes in plant reproduction and development. To guard against potential negative consequences of disrupting endogenous *S* gene functions, in the case of TALE targets, mutagenesis of EBEs rather than coding regions can be carried out, as has been demonstrated in rice and citrus^63,42^. We demonstrated the effectiveness of this approach in BBC through targeted mutagenesis of the Avrb6 EBE in the *GhSWEET10D* promoter, resulting in resistance to strain H1005. Since we have shown that the virulence contribution of Tal7b involves simultaneous activation of *GhSWEET14a*, *GhSWEET14b*, and *GhPL1*, multiplexed editing of the Tal7b EBEs in each of these genes in each subgenome in which they occur, could be a highly effective strategy for resistance to the currently prevalent *Xcm* genotypes that depend on Tal7b for virulence. This is especially true given our finding also that blocking activation of either the *SWEETs* or the *PL* gene reduces disease symptoms without impacting bacterial population growth, alleviating selection imposed on the pathogen population that would be imposed by other forms of resistance that restrict pathogen multiplication.

## MATERIALS AND METHODS

### Plant materials, bacterial strains, and growth conditions

*Gossypium hirsutum* germplasm Acala 44E (Ac44E) and FiberMax FM 2322GL (FM 2322GL) were cultivated in 3.5-inch square pots containing Metro Mix 900 soil (SunGro Horticulture, Agawam, MA, USA) in a growth chamber at 28°C, 65% relative humidity, and 100 μE m^−2^s^−1^ light with a 12-hr light/12-hr dark photoperiod. *Nicotiana benthamiana* were cultivated in 3.5-inch square pots containing Metro Mix 366 soil (SunGro Horticulture, Agawam, MA, USA) in a growth chamber at 22°C, 30% humidity and 100 μE m^−2^s^−1^ light with a 12-hr light/12-hr dark photoperiod. Cotton plants inoculated with *Xanthomonas citri* pv. *malvacearum* (*Xcm*) were transferred into a growth chamber at 28°C, 65% humidity, and 100 μE m^−2^s^−1^ light with a 12-hr light/12-hr dark photoperiod. *Escherichia coli* DH5α and *Agrobacterium tumefaciens* GV3101 were incubated in Luria-Bertani (LB) media overnight with appropriate antibiotics (ampicillin 100 µg/mL, gentamicin 20 µg/mL, kanamycin 50 µg/mL, spectinomycin 50 µg/mL, or tetracycline 10 µg/mL) on a rotary shaker at 37°C or 28°C, respectively. *Xcm* strains were incubated in Nutrient Broth (NB; 0.5% peptone, 0.3% yeast extract, 0.5% NaCl) media with appropriate antibiotics (gentamicin 20 µg/mL, kanamycin 50 µg/mL, or rifampicin 50 µg/mL) on a rotary shaker at 28°C. *Xcm* field isolates were collected from BBC-infected cotton fields in Texas by excising water-soaked lesions from infected cotton leaves with a sterile scalpel. Cut lesions were macerated in sterile water, serial diluted, and plated on Nutrient Agar (NA; 0.5% peptone, 0.3% yeast extract, 0.5% NaCl, and 1.5% agar). Individual colonies were re-streaked and inoculated into cotton to confirm pathogenicity.

### *Xcm* DNA isolation for whole-genome sequencing

Texas *Xcm* field isolates were incubated overnight in 30 mL of glucose yeast extract media (2% w/v glucose, 1% w/v yeast extract) for total DNA extraction. The bacterial suspension was centrifuged at 3000 g at 4°C, and the pellet was washed twice with sterile NE buffer (150 mM NaCl and 50 mM EDTA). The bacterial pellet was resuspended in 2.5 mL TE buffer (50 mM Tris-HCl, pH 8.0, and 50 mM EDTA) and mixed with 10 µl Ready-Lyse lysozyme (Epicentre Biotechnologies, Madison, WI, USA) and 50 µl RNase (10 mg/mL). The bacterial mix was stored at 4°C for 45 minutes before adding 1 mL STEP solution (0.5% SDS, 50 mM Tris-HCl, pH 7.5, 40 mM EDTA, and 2 mg/mL protease K) and incubating for 1 hour at 37°C. The cell lysate was neutralized with 1.8 mL 7.5 M ammonium acetate before adding 10 mL phenol:chloroform:IAA (25:24:1) and phase separated by centrifuging at 7000 g for 10 minutes. The supernatant was collected, and 10 mL chloroform:IAA was added before centrifuging at 7000 g for 10 minutes. The supernatant was collected, and 2x volume of 95% ethanol was added. The precipitated DNA was transferred to a 2 mL tube using a plastic pipette tip and centrifuged at 2000 g for 5 minutes. Excess liquid was removed, and the DNA pellet was washed with 70% ethanol. The DNA pellet was briefly air-dried and dissolved in TE buffer overnight at 4°C. The quantity and quality of DNA was checked using a NanoDrop One Microvolume UV-Vis Spectrophotometer (ThermoFisher Scientific, Waltham, MA, USA).

### PacBio whole-genome sequencing and *de novo* genome assembly of *Xcm* isolates

DNA library preparation and sequencing were conducted following the manufacturer’s instructions^64^ and performed at the Genomics Core Facility of the Icahn School of Medicine at Mount Sinai (New York, NY, USA). Electrophoresis was conducted using standard methods to detect small plasmids in each strain, which could potentially be lost during library preparation ^18^. Sequencing was conducted to >120 coverage with two Single-molecule real-time (SMRT) cells per isolate. Analysis of the read-length distribution revealed a fat tail, with approximately 20% of the coverage post-adaptor removal contained in subreads ≥15,000 bp. The whole genomes of *Xcm* TX4, TX7, and TX9 were assembled using HGAP3^65^ and validated by performing local assemblies of reads containing TALE genes. These local assemblies were then compared to the whole-genome assembly following the methodology previously described^64^.

Chromosomes and plasmids were aligned using progressiveMAUVE^66^ with default parameters, employing a progressive alignment approach. ProgressiveMAUVE was chosen due to its effectiveness in handling large-scale rearrangements, inversions, and other genomic rearrangements commonly observed in microbial genomes. Additionally, it produces alignments that uncover both conserved regions and genomic rearrangements across multiple genomes, facilitating comparative genomic and evolutionary analysis. The D-GENIES web tool^67^ with alignment by Minimap2 v2.26^68^ was used to map pair-wise comparisons of chromosomes and plasmids to generate percent identity scores. Proteins for *Xcm* TX4, TX7, TX9 were predicted using the NCBI Prokaryotic Genome Annotation Pipeline (PGAP)^69^. Phylogenetic analysis was conducted using OrthoFinder v.2.3.11^70^, which uses the STAG and STRIDE algorithms for rooting.

### Immunoblot analysis of *Xcm* TALomes

*Xcm* strains were incubated overnight in NB media on a rotary shaker at 28°C. Bacterial pellets were harvested by centrifuge and washed twice with sterile double-distilled^65^ H_2_O to remove exopolysaccharides. Bacterial cells were diluted to OD_600_ = 0.5, and 120 µl was added to 40 µl of 4x SDS buffer (250 mM Tris-HCl pH 6.8, 4% SDS, 0.1% bromophenol blue, 40% glycerol, and 4% β-mercaptoethanol) and heated to 95°C for 10 minutes. Proteins were separated in a 7.5% SDS-PAGE gel before being transferred to a PVDF membrane. PBST solution (phosphate-buffered saline solution with 0.1% Tween-20) with 5% nonfat milk was used to block the membrane for 3 hours at room temperature. TALEs were detected by immunoblotting with primary α-TALE antibodies^71^ (1:2000) in 5% milk PBST solution overnight at 4°C, followed by secondary goat-α-rabbit-IgG-HRP antibody (ThermoFisher Scientific, Waltham, MA, USA) in 5% milk PBST solution for 2 hours at room temperature. Protein bands were analyzed using a ChemiDoc Gel Imaging System (Bio-Rad, Hercules, CA, USA)

### Construction of TALE expression vectors

TALEs were individually cloned into an in-house custom Gateway entry vector *pBADZ2* using the Gibson assembly approach to *TALE* gene cloning first developed by Li et al. (2019). *pBADZ2* was digested with restriction enzymes SpeI and AvrII, while 30 µg gDNA of *Xcm* was digested with SphI at 37°C for 4 hours. Linearized *pBADZ2* was purified by 1.5% agarose gel extraction. gDNA was placed in an 80°C water bath for 15 minutes to stop the digestion. DNA was then precipitated by adding 10% of 3M potassium acetate pH 5.2 and 2x volume of 100% ethanol before centrifuging at 15,000 g for 2 minutes. The pellet was washed in 70% ethanol and centrifuged at 15,000 g for 5 minutes. The supernatant was discarded, and the DNA pellet was resuspended in sterile water. DNA quantity and quality were checked using a NanoDrop One Microvolume UV-Vis Spectrophotometer (ThermoFisher Scientific, Waltham, MA, USA). Gibson assembly was performed according to the manufacturer’s instructions using NEBuilder® HiFi DNA Assembly (New England BioLabs Inc., Ipswich, MA, USA). *E. coli* DH5a was transformed with a 5 µl reaction mix, and transformants were selected on LB plates with appropriate antibiotics (gentamicin 20 µg/mL). Thirty colonies were selected for plasmid miniprep and screened to confirm TALE inserts by digestion with restriction enzymes StuI, targeting the plasmid backbone, and AatII, targeting the insert. Gateway cloning was used to shuttle TALEs into the destination vector *pKEB1* using the manufacturer’s instructions from Gateway LR Clonase II Enzyme mix (ThermoFisher Scientific, Waltham, MA, USA). Colonies were selected for plasmid miniprep and screened by restriction enzyme digestion using SphI to confirm TALE inserts. All constructs were confirmed by Sanger sequencing.

### Construction of *Xcm* TALE mutants

Suicide vector *pTeM5* was transformed into *Xcm* TX4 by electroporation^72^. *pTeM5* contains a kanamycin resistance gene flanked by the central repeat region of AvrXa5 from *Xoo* strain PXO99A and the N- and C-termini from *Xoc* strain BLS256. Mutants were generated by single or double crossover from plasmid integration into the *Xcm* genome. The genomic DNA extracted from potential mutants was subjected to digestion with the restriction enzymes XmnI and EcoRI. These enzymes cleave DNAs at sites outside the TALE open reading frame, generating unique DNA fragments for each effector. Following digestion, the resulting DNA fragments were separated by agarose gel electrophoresis until bands were adequately separated. Southern blotting was conducted to transfer the DNA fragments from the gel onto a membrane, which was subsequently hybridized with a TALE-specific probe. The resulting Southern blot images were examined to determine the presence or absence of DNA fragments corresponding to the TALE loci in the mutant samples compared to wild-type controls. To validate TALE mutations in *Xcm*, genomic DNA digested with SphI was ligated into the *pBADZ2* vector, which had been previously digested with SpeI and AvrII, by Gibson assembly using the NEBuilder® HiFi DNA Assembly kit and following the manufacturer’s instructions (New England BioLabs Inc., Ipswich, MA, USA). The resulting Gibson assembly products were transformed into 50 µl of DH5α competent *E. coli* cells. Bacterial colonies were selected based on antibiotic marker resistance, and PCR amplicons derived from the insertion regions were subjected to Sanger sequencing. To characterize the TALE repertoire in each *Xcm* mutant, the cloned DNA sequences were translated into amino acids using the Expasy DNA translation tool (https://web.expasy.org/translate/). This translation facilitated the identification of the RVDs associated with different TALEs. Individual TALE mutants were identified by the presence of a kanamycin resistance gene flanked by RVDs corresponding to a specific TALE. *Xcm* mutants lacking specific TALEs were used for the assays.

### Bacterial tri-parental mating

Broad-host-range vector *pKEB1*, a gateway-compatible *pUFR047* derivative^21,73^, carrying TALEs, designer TALEs, or empty vector controls was transformed into *Xcm* HM2.2 by bacterial tri-parental mating. Helper plasmid *pRK2073* and modifier plasmid *pUFR054* were included to facilitate plasmid transfer from *E. coli* to *Xcm*^73^. Overnight liquid cultures of *E. coli* and *Xcm* were centrifuged at 2,000 g for 5 minutes. Bacterial pellets were washed with Nutrient broth (NB) and centrifuged again at 2,000 g for 5 minutes. Bacterial cells were individually resuspended in NB, with *Xcm* HM2.2 having a higher density compared to the other strains, in a ratio of 5:1:1:1 (recipient:donor:helper:modifier). Subsequently, 5µl of each suspension was spot inoculated on top of each other on Nutrient agar (NA) plates. NA plates were incubated overnight at 28°C. Conjugated bacteria were suspended in NB, spread onto NA plates with antibiotics, and incubated for 2 days at 28°C. Single colonies were re-streaked on NA plates with antibiotics to purify the bacteria. Conjugants were confirmed by colony PCR.

### *Xcm* inoculations and water-soaked lesion quantification

*Xcm* strains were incubated in NB liquid media with appropriate antibiotics (gentamicin 20 µg/mL, kanamycin 50 µg/mL, or rifampicin 50 µg/mL) overnight on a rotary shaker at 28°C. For syringe inoculations, bacterial suspensions were adjusted to OD_600_ = 0.05 in sterile ddH_2_O with 10 mM MgCl_2_. Small punctures were made to facilitate infiltration on the underside of expanded cotton cotyledons using a 10 µl pipette tip. Bacterial solutions were injected into plant tissue using a needleless syringe. The severity of the water-soaked lesion was quantified using ImageJ version FIJI^74^ as described^75^. For vacuum infiltration, bacterial suspensions were adjusted to OD_600_ = 0.0001 in ddH_2_O with 10 mM MgCl_2_ and 0.04% Silwet L-77 surfactant solution. Four-week-old cotton plants were submerged in the bacterial solution inside a desiccator connected to a vacuum pump. Plants were vacuum infiltrated for 5 minutes at 76 mmHg. Water-soaked lesions were manually quantified using ImageJ. The mean was calculated from three or more biological replicates.

### Cotton RNA isolation for RT-PCR and RT-qPCR analysis

Total RNA was extracted from cotton leaves using a Spectrum Plant Total RNA Kit (Sigma-Aldrich, Burlington, MA, USA) according to the manufacturer’s protocol. Extracted RNA was treated with DNase to remove residual genomic DNA, and 1 µg of purified RNA was used for reverse transcription by first-strand synthesis of cDNA with oligo(dT). Conditions for PCR amplification were 30 cycles of 15 seconds at 98°C, 30 seconds at 55°C, and 45 seconds at 72°C. *GhACTIN* was used as an internal control for RT-PCR. RT-qPCR analysis was performed using iTaq SYBR green supermix (Bio-Rad Hercules, CA, USA) with a Bio-Rad CFX384 Real-Time PCR System. Gene expression was measured in relation to *GhUBQ*^21,76^.

### *Xcm* bacterial counting

*Xcm* strains were washed and resuspended in sterile ddH_2_O with 10 mM MgCl_2_ at OD_600_ = 0.0001. Bacterial solutions were syringe inoculated into cotyledons. Leaf discs from three biological repeats were collected at 0, 1-, 3-, 5-, and 7 days post-inoculation and macerated in 100 µl of sterile ddH_2_O. Serial dilutions were plated on NA plates with appropriate antibiotics (gentamicin 20 µg/mL or kanamycin 50 µg/mL) and incubated at 28°C. Bacterial colony-forming units were counted after two days.

### Luciferase reporter assay

Reporter constructs *pGhSWEET14a::LUC* and *pGhSWEET14b::LUC* were constructed by PCR amplifying the *GhSWEET14aA* and *GhSWEET14bD* promoters from gDNA of Ac44E using primers containing BamHI and NcoI restriction sites. Promoter sequences were designated as 1000 bp upstream from the translational start site. PCR amplicons were cloned into binary vector *pCB302* immediately upstream on a luciferase gene^77^. Reporter constructs were transformed into *A. tumefaciens* GV3101 using electroporation. The leaves of five-week-old *N. benthamiana* were syringe-infiltrated with *Agrobacterium* containing reporter constructs and resuspended in infiltration buffer (10 mM MES, pH 5.7, 10 mM MgCl_2_, and 200 µl acetosyringone) at OD_600_ = 1.5. Inoculated *N. benthamiana* were covered with a plastic dome and placed under ambient light for 24 hours before co-infiltrated with *Xcm* strains resuspended in 10 mM MgCl_2_ at OD_600_ = 0.5. Leaf discs from co-infiltrated tissue were harvested 24 hours post-inoculation and evenly sprayed with 0.2 mM luciferin with 0.2% Silwet L-77. Luciferase activity was detected by Glomax Multi-Detection System (Promega^TM^ Madison, WI, USA).

### Virus-induced gene silencing

A conserved region between *GhSWEET14aA*, *GhSWEET14aD*, and *GhSWEET14bD*, or *GhPL1* was PCR amplified using primers containing EcoRI and KpnI restriction sites and cDNA from *G. hirsutum*. PCR amplicons were cloned into modified *Tobacco rattle virus* vector *pYL156* (*pTRV-RNA2*). *A. tumefaciens* containing *pYL156-GhSWEET14a/b*, *pYL156-GhPL1*, *pYL156*-empty vector, or *pYL192* (*pTRV-RNA1*) were resuspended in infiltration buffer (10 mM MES, pH 5.7, 10 mM MgCl_2_, and 200 µl acetosyringone) at OD_600_ = 1.5. Bacterial suspensions of *pYL192* were mixed with *pYL156-GhSWEET14a/b, pYL156-GhPL1,* or *pYL156*-empty vector at a 1:1 ratio before syringe inoculating into cotton cotyledons. Inoculated cotton plants were covered with a plastic dome and placed under ambient light overnight, then moved to a growth chamber at 22°C, 30% humidity, and 100 μE m^−2^s^−1^ light with a 12-hr light/12-hr dark photoperiod^78^.

### Construction of designer TALEs

Designer TALEs were constructed using the Golden Gate TALEN and TALE Kit 2.0 (Addgene, Watertown, MA, USA) according to the manufacturer’s protocol^79^. EBE target sequences were evaluated for dTALE binding efficiency using the TAL Target Finder tool^38^. The central repeat region of each dTALE consisted of repeats containing RVD sequences corresponding with the respective nucleotide. dTALEs were shuttled into broad-host-range vector *pKEB1* using the Gateway LR Clonase II Enzyme mix (ThermoFisher Scientific, Waltham, MA, USA) according to the manufacturer’s instructions.

### Construction of CRISPR-Cas9 vectors

Single-stranded oligonucleotides were synthesized comprising of the guide RNA (gRNA) target sequence for *pGhSWEET10D* (5’-AGAACGGGCTGGGGATG-3’) with a 5’-CGGA motif on the forward oligonucleotide and a 5’-AAAC motif on the reverse oligonucleotide. The oligonucleotides were annealed together and cloned into the BsaI-linearized *pTX171* vector using the Golden Gate method as previously described^80^.

For mutating *GhSWEET14a* and *GhSWEET14b,* single-stranded oligonucleotides were designed according to established protocols^40^. The *pGTR* plasmid, containing a fused tRNA-gRNA scaffold fragment, served as the DNA template for PCR to construct a polycistronic tRNA-gRNA (PTG) gene encompassing the gRNA target sequences for *GhSWEET14a* (5’-ATATACCAAAAACAACAGCCA-3’) and *GhSWEET14b* (5’-AGACACCAAATGCAACTGCCA-3’). Subsequently, the PTG gRNA-GhSWEET14a/b product was cloned into the *pRGEB32-GhU6.9* vector using the Golden Gate method.

All gRNA target sequences were generated using the gRNA design tool CRISPR-P (http://cbi.hzau.edu.cn/crispr). The resulting Golden Gate ligation products were transformed into 50 μl of competent *E. coli* DH5α cells. Bacterial colonies were selected based on antibiotic marker resistance, and vectors were validated by Sanger sequencing of PCR amplicons derived from the insertion regions.

### Generation of transgenic cotton

*Agrobacterium*-mediated cotton variety Coker 312 transformation was performed at Texas A&M Multicrop Transformation Facility following a previously described protocol with modifications^81^. Briefly, cotton seedlings were cultivated using Murashige and Skoog (MS) medium. Hypocotyls were excised and infected with *A. tumefaciens* LBA4404 containing the CRISPR-Cas9 construct or a GFP control. Explants and *Agrobacterium* were co-cultivated on P1 [4.31 g/L MS salts, 100 mg/L myoinositol, 0.4 mg/L thiamine HCl, 5 mg/L *N*^6^-(2-isopentenyl)adenine, 0.1 mg/L α-naphthaleneacetic acid (NAA), 3% glucose, 1 g/L magnesium chloride hexahydrate, pH 5.8, 0.2% Phytagel] medium supplemented with 50 µM acetosyringone for 72 hours under light at 25°C before being transferred to P1 media containing 15 mg/L hygromycin. Explants were cultivated in a growth chamber at 28°C for 42 days until callus tissue developed at the cut surface of explants. Callus tissue was excised under a microscope, transferred to P1 media containing 15 mg/L hygromycin, and incubated under light at 28°C. Somatic embryos were transferred to EG3 (2.16 g/L MS salts, 0.5% glucose, 100 mg/L myoinositol, 0.4 mg/L thiamine HCL, 0.01 mg/L NAA, pH 6.5, 0.2% Phytagel) media for regeneration and subsequent plantlets were transferred to MS3 (2.16 g/L MS salt, 0.5% glucose, 0.14 mg/L thiamine HCl, 0.1 mg/L pyridoxine HCl, 0.1 mg/L nicotinic acid, pH 5.8, 0.08% Phytagel, 0.6% Bacto agar) media for further growth and development before being transferred to soil. Genomic DNA was extracted from the edited line, and the target site was amplified by PCR, and the amplicon sequenced to identify homozygous lines. GFP lines were further confirmed by confocal microscopy observation.

### RNA sequencing and transcriptomic profiling

Two-week-old cotyledons of cotton variety FM 2322GL were syringe-inoculated with *Xcm* HM2.2 expressing Tal7b or a vector control. Total RNA was extracted 24 hours post-inoculation (hpi) from three independent biological repeats. RNA-seq libraries were prepared using Illumina TruSeq Stranded mRNA Sample Preparation Kit following the manufacturer’s protocol and sequenced using an Illumina HiSeq 2000 platform with 2×150-nucleotide pair-end reads at the Texas A&M Institute for Genome Sciences and Society (College Station, TX, USA). RNA-seq analysis was performed as previously described^82^. Briefly, the Trimmomatic tool^83^ was used to preprocess raw RNA-seq reads by performing quality filtering and trimming of low-quality bases from sequencing reads to ensure that only high-quality reads were used for alignment and assembly. The alignment of RNA-seq reads to the *G. hirsutum* (AD1) TM-1 genome^29^ was executed using HISAT2^84^, employing an indexing scheme based on the Burrows-Wheeler transform and the Ferragina-Manzini index. Transcriptional assembly was conducted using StringTie^85^, which reconstructed full-length transcripts from aligned reads while estimating their abundance and quantifying gene expression levels. Differential gene expression was analyzed using Cuffdiff^86^, a component of the Cufflinks suite, which facilitated the comparison of transcript abundances between different treatments. Genes were considered differentially expressed if they demonstrated a log_2_ fold expression change >1 and an adjusted *p*-value < 0.05. These cutoffs were employed to ensure that only genes displaying statistically significant changes in expression between treatments were included in the analysis.

### EBE predictions

TALE-NT 2.0 Target Finder was used to predict TALE EBE sites within the promoterome of *G. hirsutum*^38^. Promoter regions were defined as 1000 bp upstream of the transcriptional start site. Promoter sequences were identified from the *G. hirsutum* (AD1) TM-1 genome assembled using SOAPdenovo12^29^. Binding sites passing the Target Finder score ratio cutoff of 4.0 were ranked based on Target Finder output and genomic context using a previously described machine learning classifier^54,87^.

All the primers used in this study are in **Supplementary Table 4**.

### GenBank accession numbers

*Xcm* H1005: GenBank accession numbers – CP013004 (chromosome), CP013005 (*pXcmH*), NCBI BioSample – SAMN04166563

*Xcm* N1003: GenBank accession numbers CP013006 (chromosome), CP013007 (*pXcmN*), NCBI BioSample – SAMN04166615

MSCT1: GenBank accession numbers – CP017020 (chromosome), CP017021 (*pMSCT15kb*, CP017022 (*pMSCT44kb*), CP017023 (*pMSCT60kb*), NCBI BioSample – SAMN05595756

*Xcm* MS14003: GenBank accession numbers – CP023159 (chromosome), CP023160 (unnamed plasmid 1), CP023161 (unnamed plasmid 2), CP023162 (unnamed plasmid 3), NCBI BioSample – SAMN07447517

*Xcm* AR81009: GenBank accession numbers – CP023155 (chromosome), CP023156 (unnamed plasmid 1), CP023157 (unnamed plasmid 2), CP023158 (unnamed plasmid 3), NCBI BioSample – SAMN07447516

*Xcm* HD-1: GenBank accession numbers – CP046019 (chromosome), CP046020 (unnamed plasmid 1), CP046021 (unnamed plasmid 2), NCBI BioSample – SAMN13193097

*Xcm* TX4: GenBank accession number – CP157605 (chromosome_3), CP157606 (*pXCM15_3*), CP157607 (*pXCM52.4_3*), CP157608 (*pXCM52_3*), NCBI BioSample – SAMN41592776

*Xcm* TX7: GenBank accession number – CP157601 (chromosome_2), CP157602 (*pXCM15_2*), CP157603 (*pXCM52.4_2*), CP157604 (*pXCM52_2*), NCBI BioSample – SAMN41592777

*Xcm* TX9: GenBank accession number – CP157597 (chromosome_1), CP157598 (*pXCM15_1*), CP1575599 (*pXCM52.4_1*), CP157600 (*pXCM52_1*), NCBI BioSample – SAMN41592778

### Quantification and statistical analysis

Data for quantification analyses are presented as mean ± standard error (s.e.) or standard deviation (s.d.) as indicated in the figure legends. The statistical analyses were performed by unpaired two-tailed Student’s *t*-test. The number of biologically independent replicates is shown in the figure legends. The *p*-values are provided in the graphs.

---

**Supplementary Figure 1.**
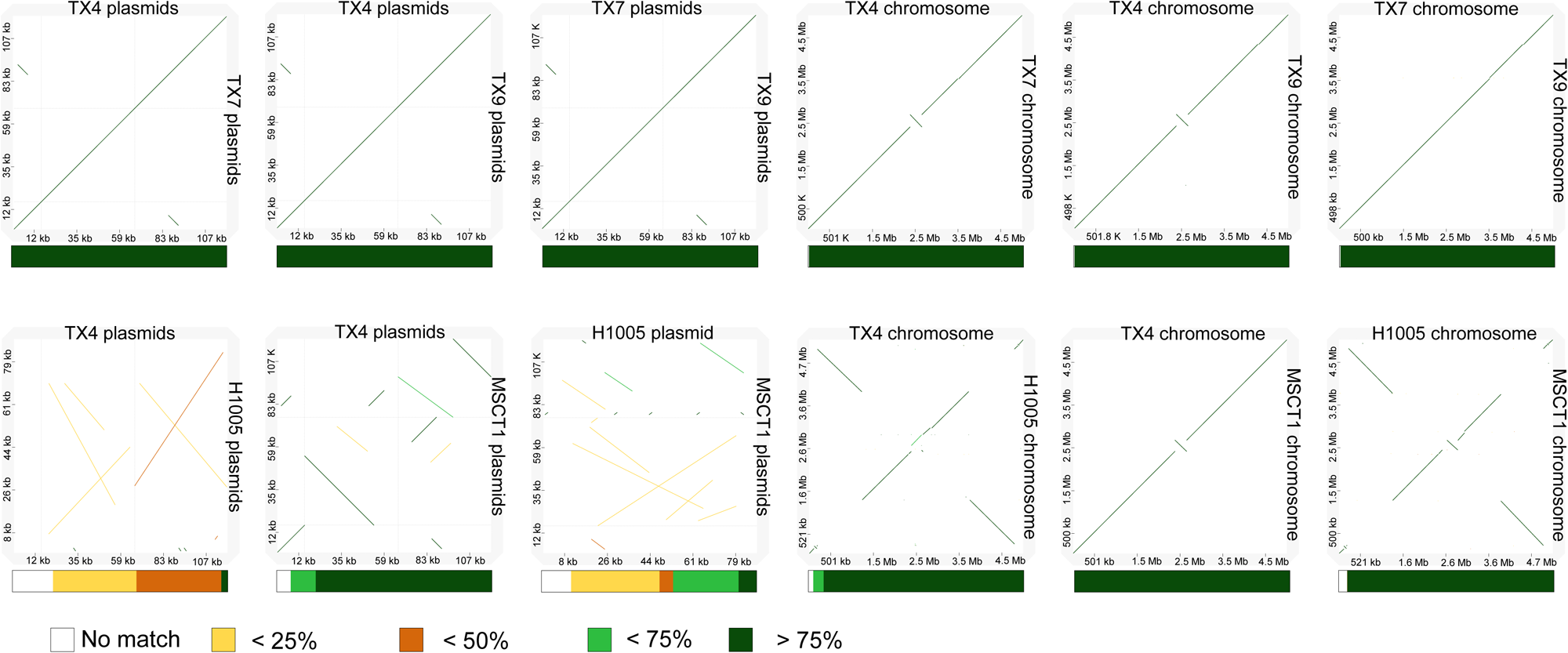
Pair-wise comparison dot plots of *Xcm* isolates with percent identity summaries. Dot plot comparisons were conducted using the Dot plot large Genomes in an interactive, efficient, and simple way (D-GENIES) tool^67^, with alignment performed by Minimap2 v2.26^68^. This approach facilitated the mapping of pairwise comparisons of both plasmids and chromosomes among isolates, enabling the generation of percent identity scores. Notably, plasmids and chromosomes were analyzed separately due to their significant scale differences, ensuring an accurate and detailed assessment of genomic relationships. The colored bars indicate percent identity scores, labeled at the bottom of each comparison.

**Supplementary Figure 2.**
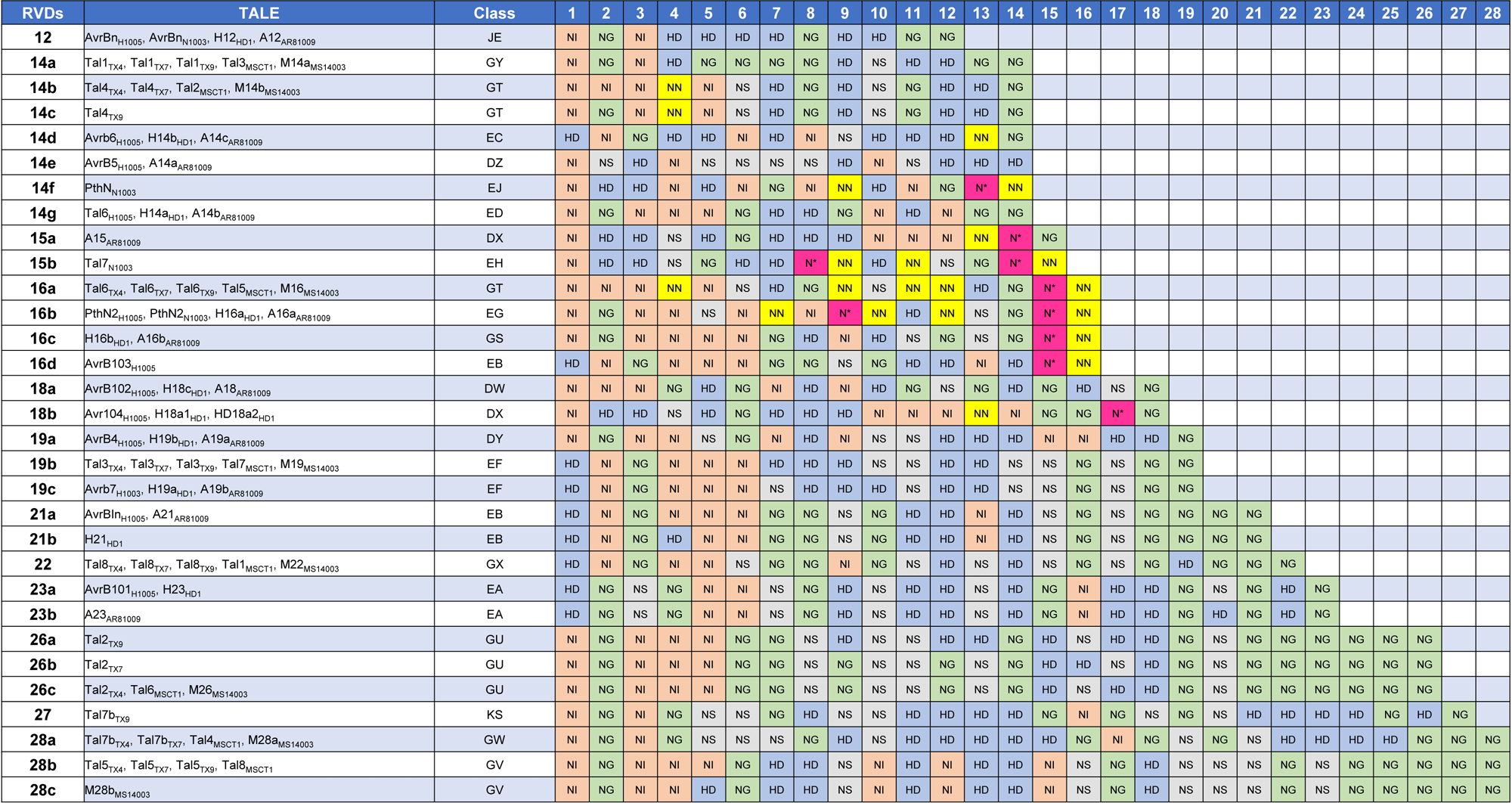
The thirty-one unique repeat variable di-residue (RVD) sequences represented among the TALEs of the nine *Xcm* strains examined. Thirty-one unique RVD sequences were identified from eighty TALEs, utilizing AnnoTALE^31^ analysis of full genome sequences obtained from nine *Xcm* isolates. The number of RVDs within the central repeat region of each TALE is indicated in the left column. A letter following the RVD number denotes separate TALEs exhibiting different RVD sequences but the same number of RVDs. In the second column, each TALE is paired with its respective *Xcm* strain. TALEs are grouped based on shared RVD sequences, with each row representing a distinct arrangement of RVDs. The numerical values in the top row represent the position of the RVD within the central repeat region. Colored boxes contain the two amino acids at of each RVD, using the single letter code. An asterisk (*) denotes a missing amino acid residue within the RVD.

**Supplementary Figure 3.**
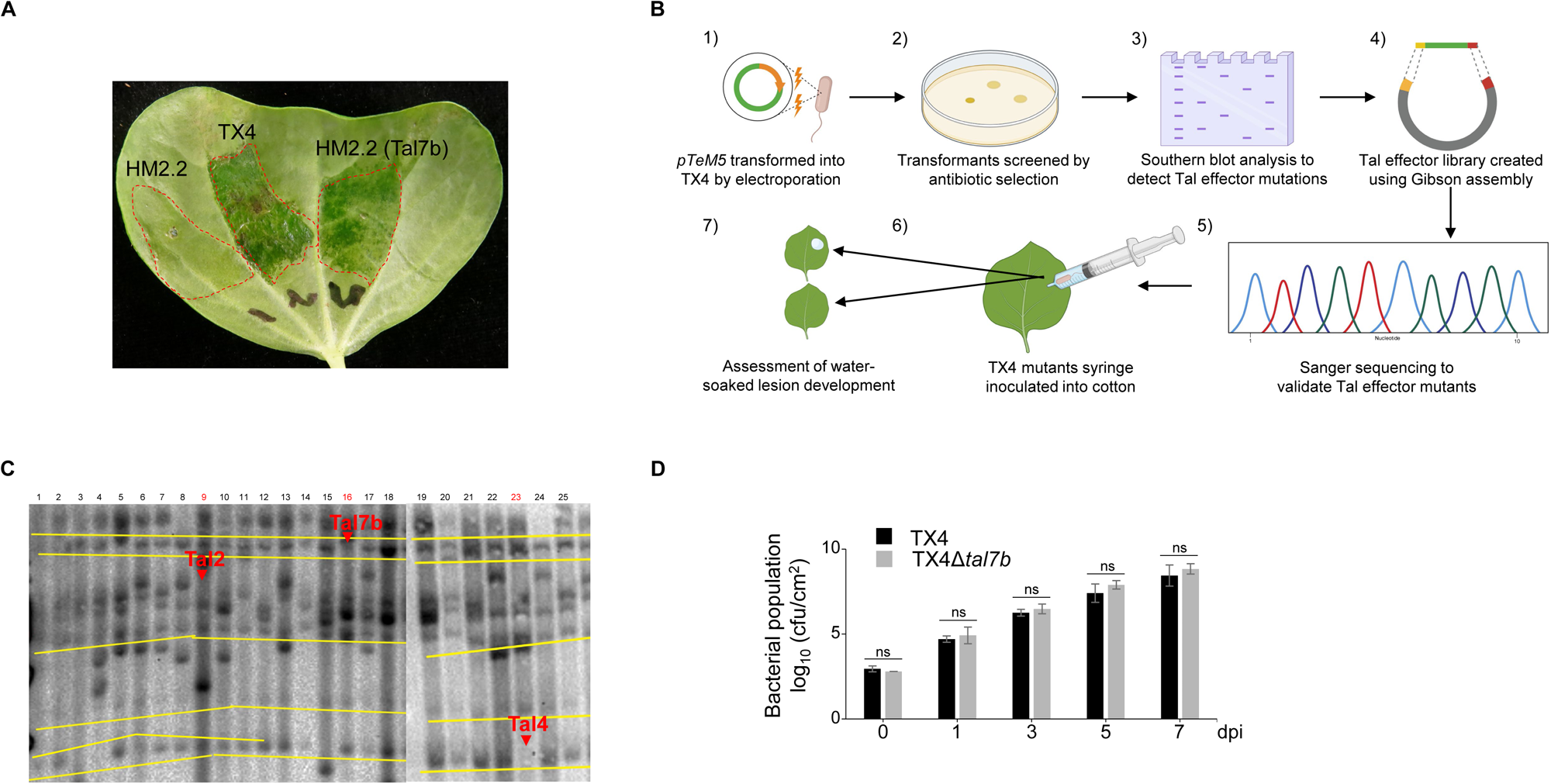
Effects of Tal7b on *Xcm* TX4 virulence. **(A)** *Xcm* HM2.2 carrying Tal7b induces water-soaked symptoms to a degree comparable to *Xcm* TX4. Different bacterial suspensions were syringe-inoculated into ten-day-old cotton cotyledons at OD_600_= 0.1. Photos were taken at 9 dpi. **(B)** Schematic illustration of generating *Xcm* TX4 TALE knockout mutants. The figure was made by BioRender. **(C)** Southern blot analysis verifies the knockout of TALE genes from *Xcm* TX4 mutants. The genomic DNA of the putative knockout mutants was digested with restriction enzymes XmnI and EcoRI for Southern blot analysis. 1 to 25 are individual mutants. Red arrows denote the expected locations of the TALEs. The absence of a band indicates the knockout of the corresponding TALE gene. **(D)** The deletion of *Tal7b* does not affect the *planta* multiplication of *Xcm* TX4. Ten-day-old cotton cotyledons were syringe-inoculated with bacterial solutions at OD_600_ = 0.0001. Leaf discs were collected at indicated time points. Data are shown as mean ± s.d. (*n* = 3) from three independent repeats. There was no significant difference between *Xcm* TX4Δ*tal7b* and TX4 using a two-tailed Student’s *t*-test.

**Supplementary Figure 4.**
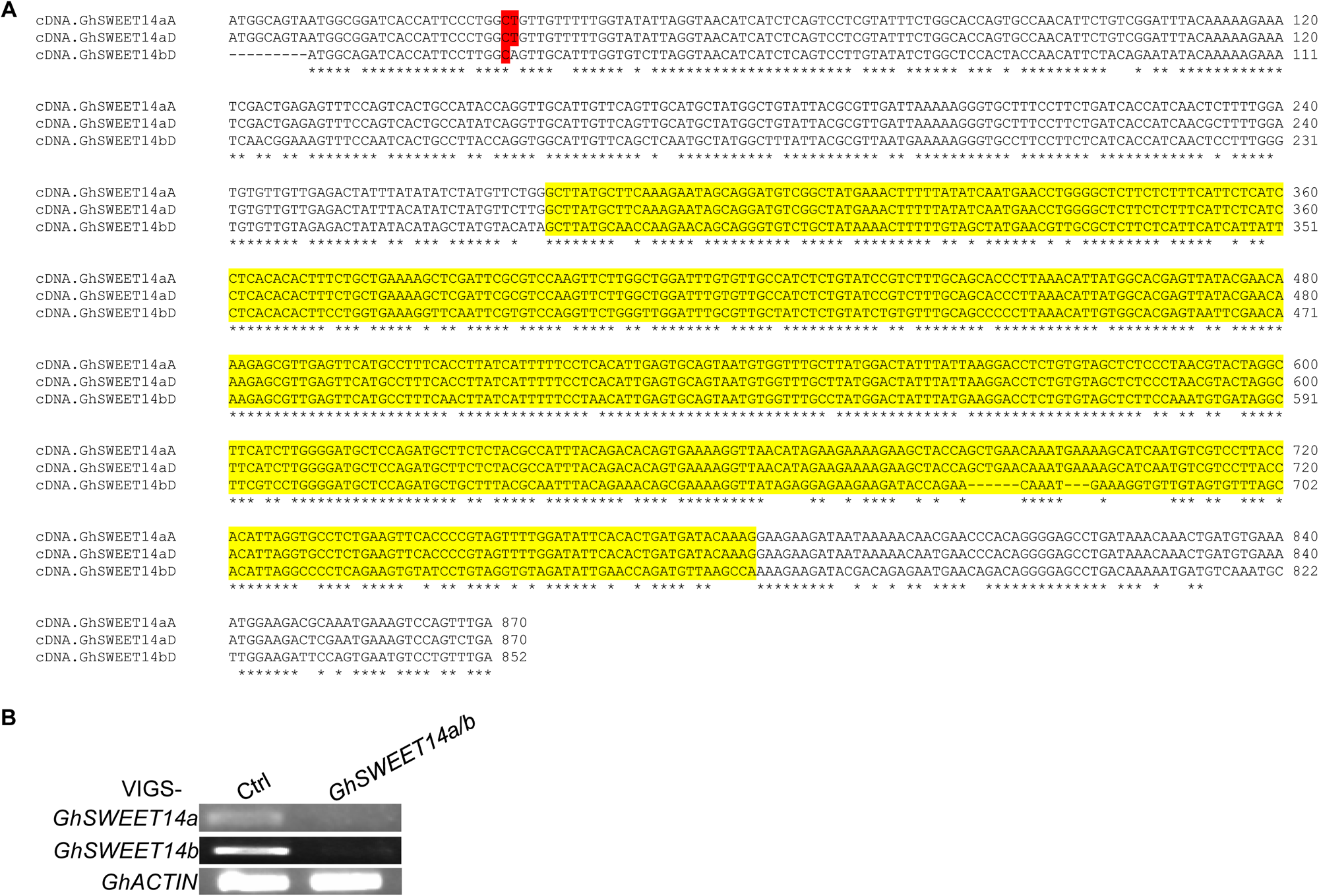
Gene sequences of *GhSWEET14a* and *GhSWEET14b* and VIGS efficiency. **(A)** Complementary DNA (cDNA) sequences of *GhSWEET14aA*, *GhSWEET14aD*, and *GhSWEET14bD*. Gene sequences were obtained from the CottonGen database^29,88^ and aligned using Clustal Omega Multiple Sequence Alignment^89^. The yellow highlighted region indicates the conserved sequence targeted for virus-induced gene sequencing (VIGS) of *GhSWEET14aA, GhSWEET14aD,* and *GhSWEET14bD.* Red highlighted regions indicate CRISPR-induced deleted nucleotides. **(B)** The basal expression of *GhSWEET14a* and *GhSWEET14b* is reduced upon virus-induced gene silencing. Seven-day-old cotton cotyledons were syringe-inoculated with *Agrobacterium* carrying VIGS constructs to silence both *GhSWEET14a* and *GhSWEET14b* or vector control. Three weeks later, the 2^nd^ true leaves were harvested for RNA extraction and RT-PCR analysis of *GhSWEET14a* and *GhSWEET14b* expression. *GhACTIN* was used as an internal control.

**Supplementary Figure 5.**
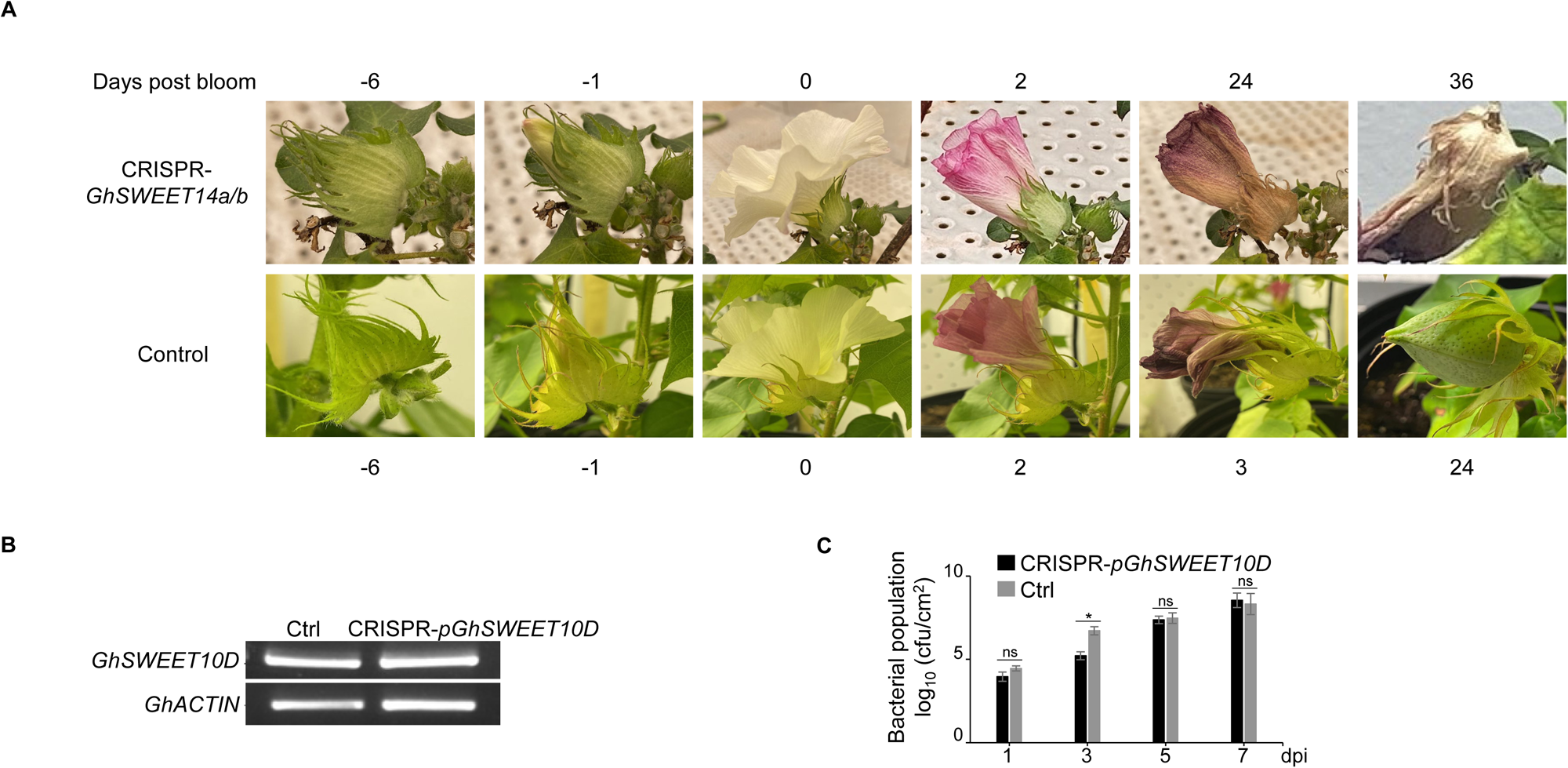
CRISPR-mediated gene-editing of *GhSWEETs* in cotton boll development and BBC resistance. **(A)** CRISPR-mediated editing of *GhSWEET14a/b* impairs boll development in cotton. The control plants exhibited typical flower withering and subsequent boll formation a few days after bloom, whereas the flowers of the CRISPR-*GhSWEET14a/b* plant persisted for over 20 days without any observable boll development. **(B)** Basal expression of *GhSWSEET10D* is not changed in CRISPR-*pGhSWEET10D* cotton. Ten-day-old transgenic cotton cotyledons were harvested for RNA extraction and RT-PCR analysis of *GhSWEET10D* expression. *GhACTIN* was used as an internal control and PCR amplification was performed for 30 cycles. **(C)** *Xcm* H1005-mediated induction of *GhSWEET10D* does not affect bacterial multiplication *in planta* following cotton infection. Ten-day-old transgenic cotton cotyledons were syringe-inoculated with *Xcm* H1005 at OD_600_ = 0.0001. Leaf discs were collected at indicated time points. The data is shown as mean ± s.d. (*n* = 3) from three independent repeats. There was no significant difference after three days between CRISPR-*pGhSWEET10D* cotton and controls using a two-tailed Student’s *t*-test.

**Supplementary Figure 6.**
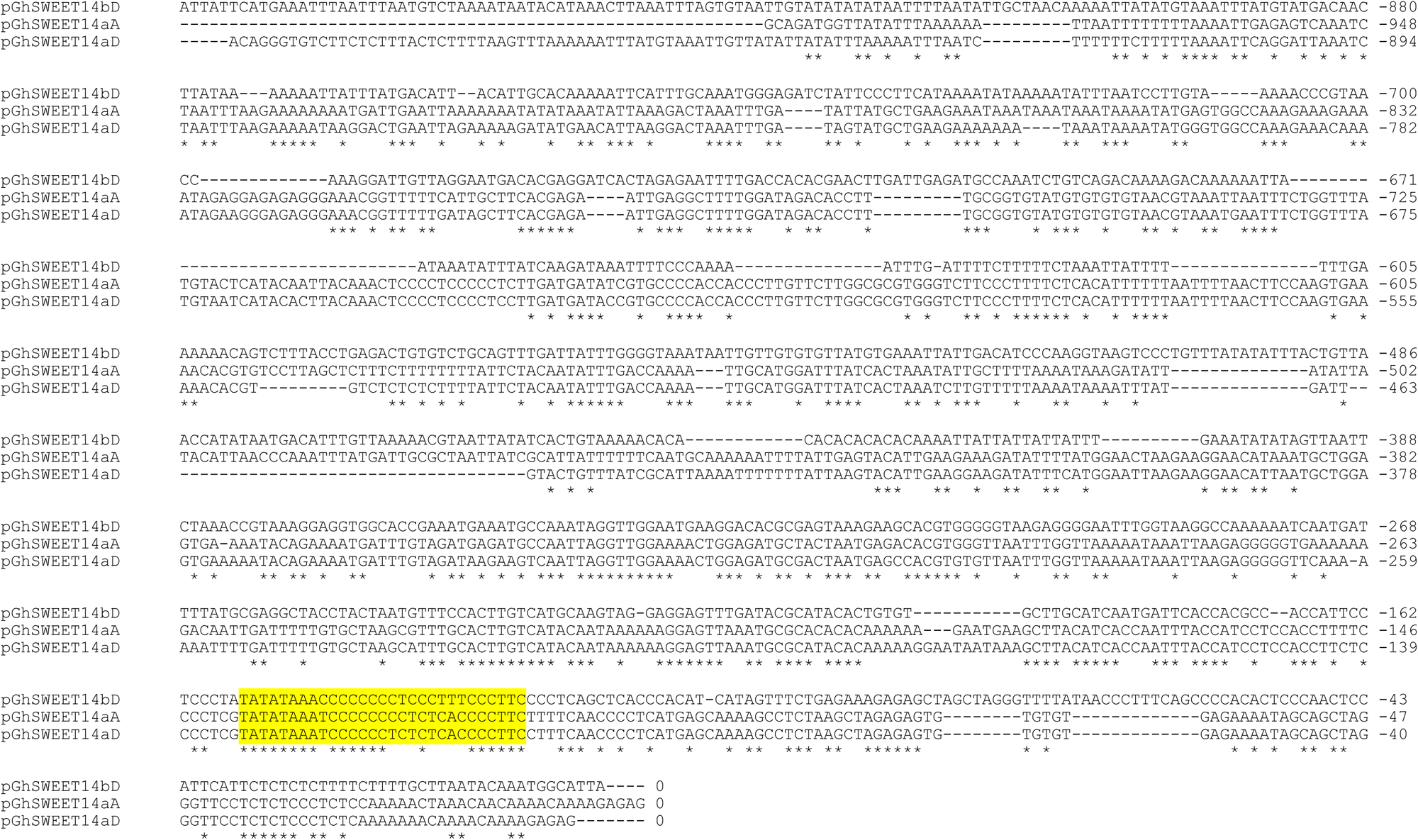
Promoter sequences of *GhSWEET14aA, GhSWEET14aD,* and *GhSWEET14bD*. Sequences were obtained from the CottonGen database^29,88^ and promoter sequences were designated as 1000 bp upstream from the translational start site. Sequence alignments were performed using Clustal Omega Multiple Sequence Alignment^89^. Highlighted sequences indicate EBE sites for Tal7b, beginning with the T at position 0^38^.

**Supplementary Figure 7.**
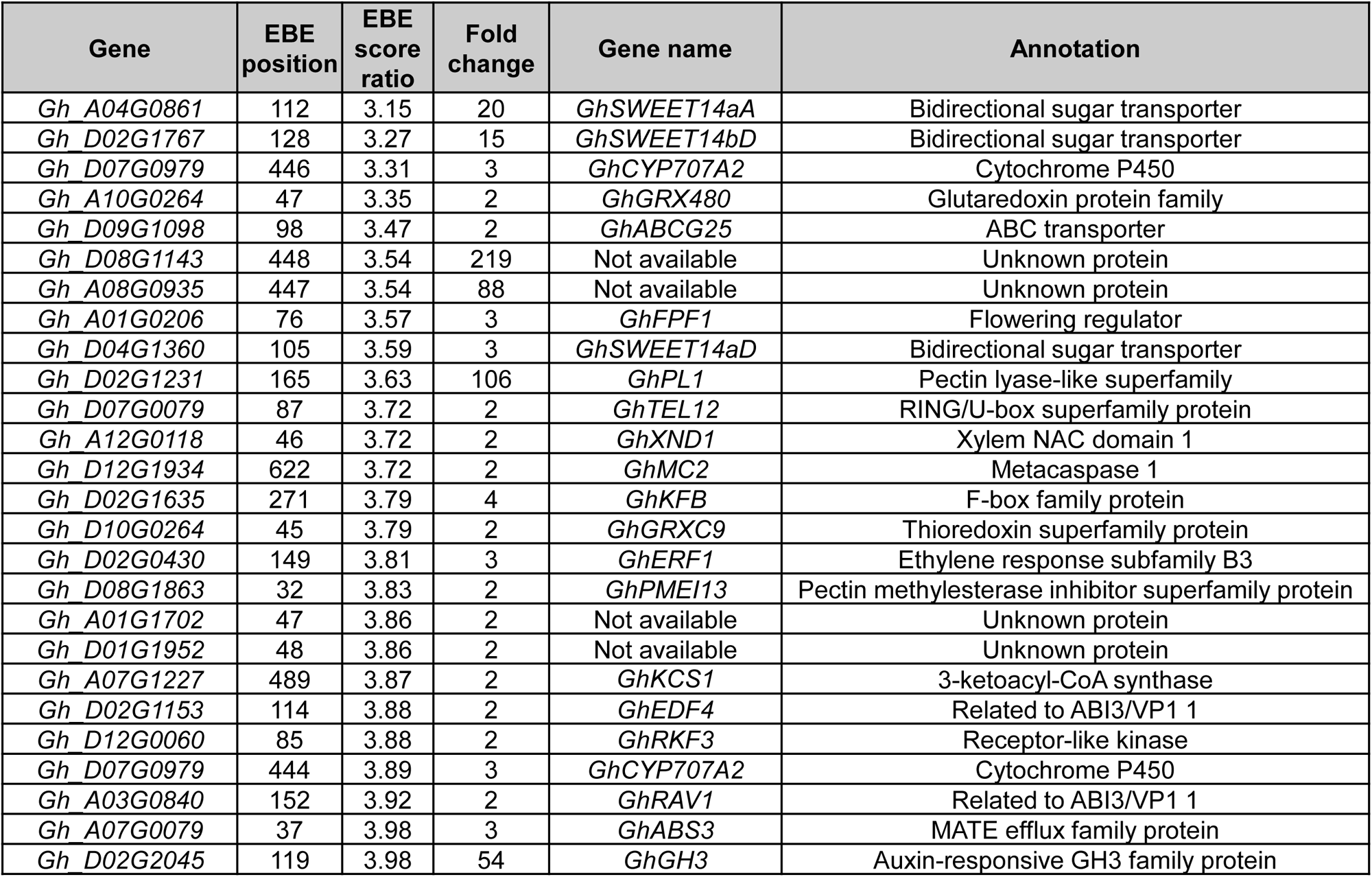
Twenty-six candidate gene targets of Tal7b. RNA-seq analysis identified 707 genes induced by *Xcm* HM2.2 Tal7b compared to *Xcm* HM2.2 using cutoffs of log_2_ fold change > 1 and *p*-value < 0.05. Cross-referencing these 707 Tal7b-induced genes with the 397 genes containing predicted Tal7b EBE sites within their promoters using a score ratio cutoff of 4.0 identified 26 candidate gene targets of Tal7b, which were ranked by the strength of the EBE score ratio and promoter context of the EBE^38^. EBE position indicates its proximity to the translational start codon.

**Supplementary Table 1.** Statistical summary of whole-genome sequencing for Texas *Xcm* isolates. The reemerging Texas *Xcm* genomes were sequenced using the long-read, Pacific Biosciences (PacBio) single-molecule real-time (SMRT) sequencing platform and assembled *de novo*. Assembled genomes have a coverage depth ranging from 193-891x with mapped subread n50 lengths of 12.9-14.9 kb.

| Isolate | Mapped subread<br>n50 | Replicon | Replicon length<br>(bp) | Coverage depth |
| --- | --- | --- | --- | --- |
| TX4 | 12,910 | Chromosome | 5,018,325 | 891x |
|  |  | Plasmid 1 | 15,295 | 261x |
|  |  | Plasmid 2 | 52,393 | 829x |
|  |  | Plasmid 3 | 52,085 | 761x |
| TX7 | 13,413 | Chromosome | 5,006,648 | 723x |
|  |  | Plasmid 1 | 15,283 | 193x |
|  |  | Plasmid 2 | 52,393 | 470x |
|  |  | Plasmid 3 | 52,084 | 590x |
| TX9 | 14,880 | Chromosome | 4,980,308 | 812x |
|  |  | Plasmid 1 | 15,283 | 210x |
|  |  | Plasmid 2 | 52,392 | 362x |
|  |  | Plasmid 3 | 51,983 | 440x |

**Supplementary Table 2.** BUSCO analysis is used to assess the completeness and quality of genome assemblies. Genome completeness was evaluated using Benchmarking Universal Single-Copy Orthologs v5.1.2 (BUSCO)^90^. All TX isolate genomes cover 100% of the genes in the bacteria_obd10 lineage and over 99% of the genes in the xanthomonadales_obd10 lineage.

| Isolate | TX4 | TX7 | TX9 |
| --- | --- | --- | --- |
| BUSCO: bacteria_obd10 (%) |  |  |  |
| Complete (C) | 100 | 100 | 100 |
| Complete and single-copy (S) | 100 | 100 | 100 |
| Complete and duplicated (D) | 0 | 0 | 0 |
| Fragmented (F) | 0 | 0 | 0 |
| Missing (M) | 0 | 0 | 0 |
| Total groups search (n) | 124 | 124 | 124 |
| BUSCO: xanthomonadales_odb10 (%) |  |  |  |
| Complete (C) | 99.6 | 99.5 | 99.5 |
| Complete and single-copy (S) | 99.6 | 99.5 | 99.5 |
| Complete and duplicated (D) | 0 | 0 | 0 |
| Fragmented (F) | 0.1 | 0.2 | 0.2 |
| Missing (M) | 0.3 | 0.3 | 0.3 |
| Total groups search (n) | 1152 | 1152 | 1152 |

**Supplementary Table 3.** *Xcm* strains used for TALome analysis. Nine *Xcm* isolates, comprising three newly sequenced Texas isolates collected during this study, and an additional six isolates whose whole genome sequences, genomic properties, and sequencing platforms were retrieved from the NCBI database.

| Isolate | Isolate origin | Genome size (bp) | Chromosome size (bp) | Plasmid number/size | Sequencing platform | TALome | Xoplome | Virulence | Reference |
| --- | --- | --- | --- | --- | --- | --- | --- | --- | --- |
| TX4 | Plains, TX, USA | 5,138,098 | 5,018,325 | 3; 52kb, 52kb, 15kb | SMRT; Minlon | 8 | 27 | Moderate | This study |
| TX7 | Matagorda, TX, USA | 5,126,408 | 5,006,648 | 3; 52kb, 52kb, 15kb | SMRT; Minlon | 8 | 27 | Moderate | This study |
| TX9 | Seminole, TX, USA | 5,099,966 | 4,980,308 | 3; 52kb, 52kb, 15kb | SMRT; Minlon | 8 | 27 | Weak | This study |
| MSCT1 | Yazoo City, MS, USA | 5,123,943 | 5,004,182 | 3; 60kb, 44kb, 15kb | SMRT | 8 | 27 | Moderate | Showmaker et al. 2017 |
| MS14003 | Wilzone, MS, USA | 5,144,706 | 5,029,617 | 3; 52kb, 47kb, 15kb | SMRT | 8 | 27 | Moderate | Phillips et al. 2017 |
| H1005 | Stillwater, OK, USA | 5,300,847 | 5,212,564 | 1; 88kb bp | SMRT; Minlon | 12 | 26 | High | Cox et al. 2017 |
| N1003 | Burkina Faso, Africa | 5,278,251 | 5,218,607 | 1; 59kb | SMRT | 4 | 26 | Weak | Cox et al. 2017 |
| AR81009 | Argentina | 5,408,848 | 5,267,057 | 3; 92kb, 26kb, 22kb | SMRT | 12 | 26 | High | Phillips et al. 2017 |
| HD-1 | China | 5,341,921 | 5,255,091 | 2; 43kb, 43kb | SMRT | 12 | 27 | N/A | N/A |

**Supplementary Table 4.**
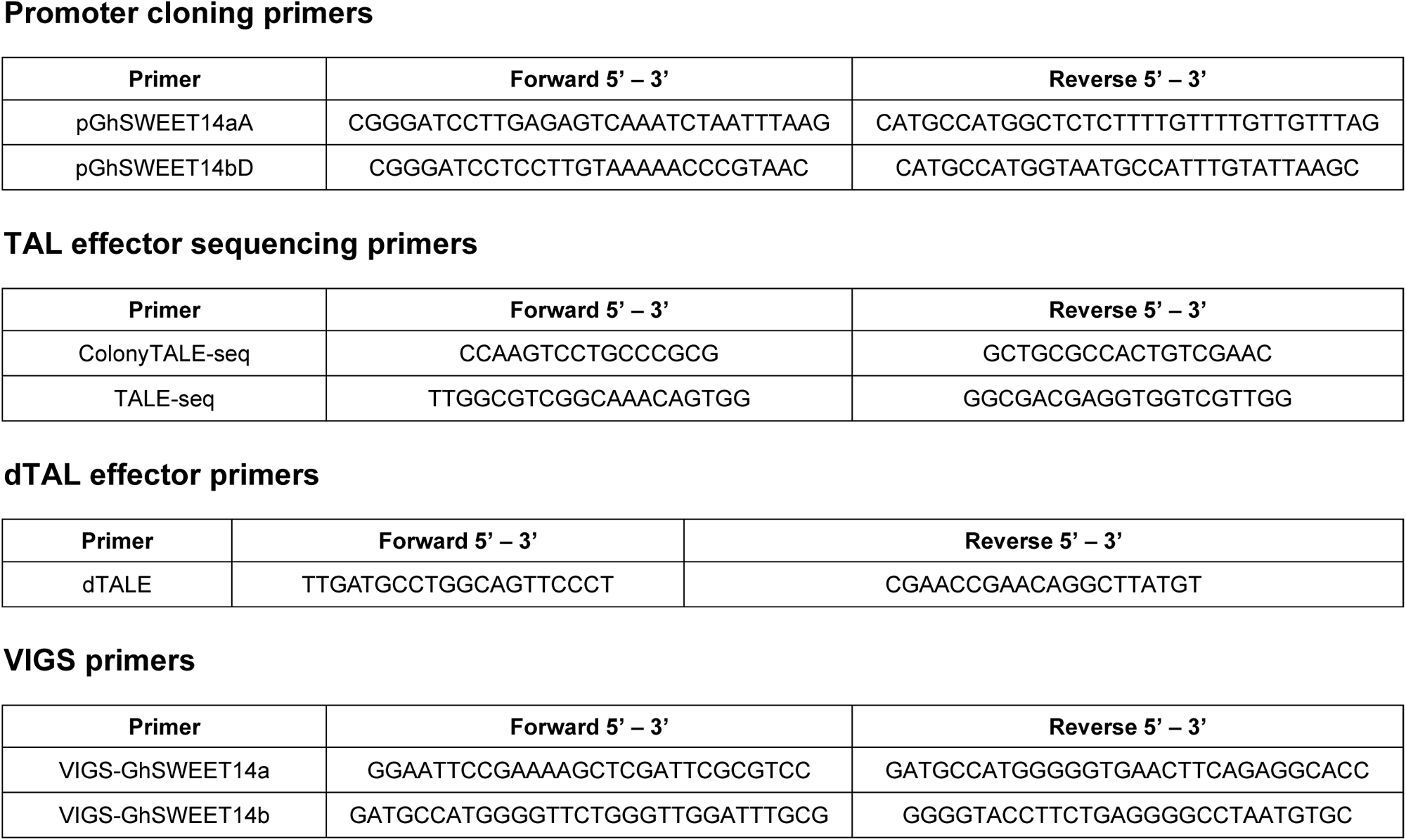
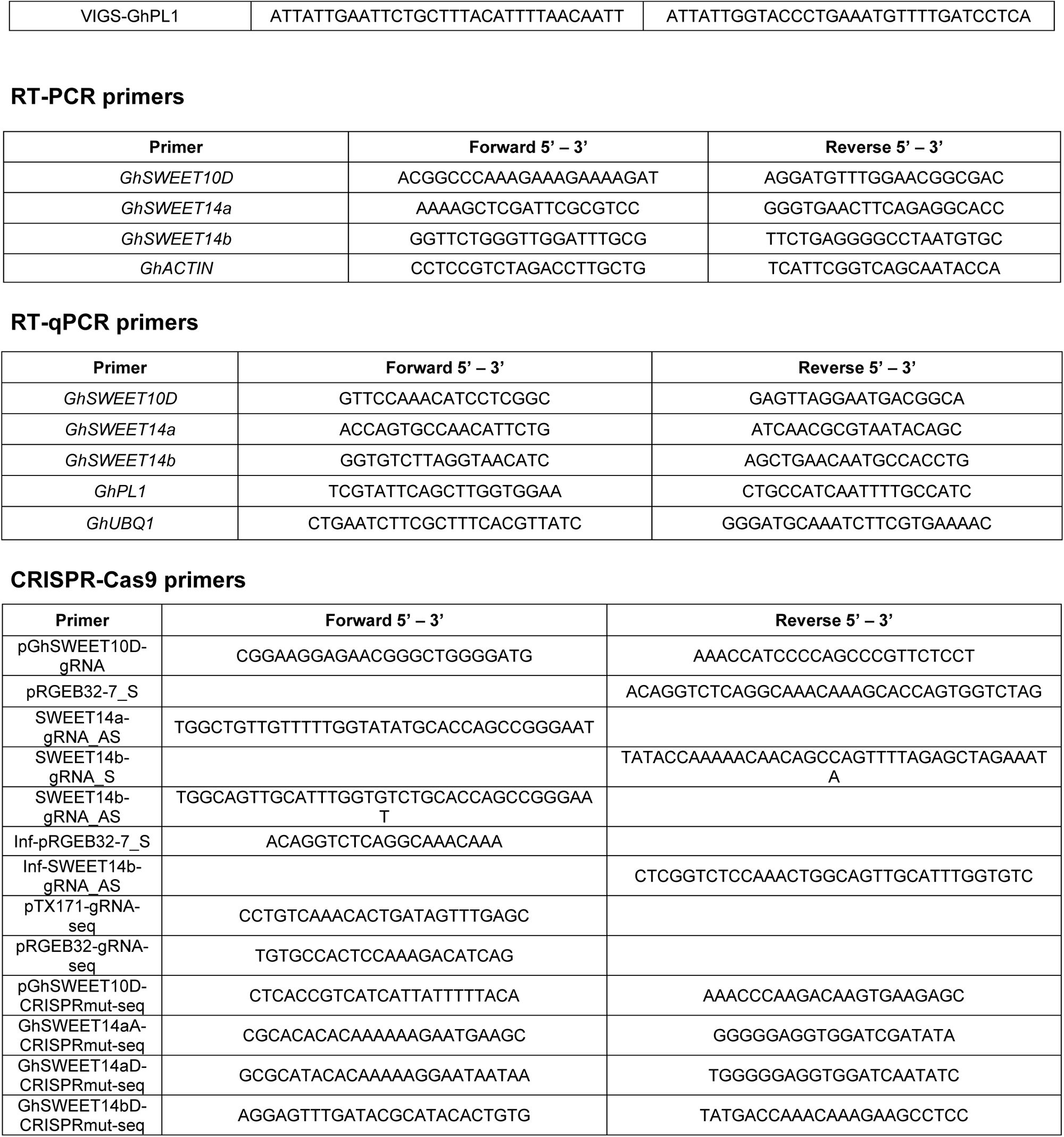
Primers used in this study.

## ACKNOWLEDGMENTS

We thank Baden Aniline and Soda Factory (BASF) for providing FiberMax cotton seeds; Dr. Shi-En Lu (Mississippi State University) for *Xcm* strain MSCT1; Leslie Wells (Texas A&M AgriLife Research) for cotton seed propagation; and members of the laboratories of L.S. and P.H. for discussions and comments on the experiments. The study was supported by the United States Department of Agriculture-National Institute of Food and Agriculture (USDA-NIFA) (2018-67013-28513) to L.S and A.J.B., and Cotton Incorporated (18-123TX) to L.S.

## AUTHOR CONTRIBUTIONS

B.M., P.H., A.J.B., and L.S. conceived the project, designed experiments, and analyzed data. B.M. performed most of the bacterial inoculation, molecular cloning, gene expression, and RNA-seq experiments. T.B and L.W. generated the *Xcm* TX4 mutants and designer TALEs. R.R. and S.C.D.C. performed genome assembly and analysis of Texas *Xcm* isolates. C.D.C. performed transcriptome analysis of RNA-seq data. K.C. initially performed RT-PCR analysis and constructed VIGS-*SWEET14a/b* vector. L.Z. constructed the CRISPR vectors. X.M. performed TALomes immunoblot analysis. T.A.W. and J.K.D. collected *Xcm* isolates from Texas cotton fields. B.M., P.H., A.J.B., and L.S. wrote the manuscript with input from all authors.

## Notes

### Competing Interest Statement

The authors have declared no competing interest.

